# Multi-omic profiling of histone variant H3.3 lysine 27 methylation reveals a distinct role from canonical H3 in stem cell differentiation

**DOI:** 10.1101/2021.08.31.458429

**Authors:** Yekaterina Kori, Peder J. Lund, Matteo Trovato, Simone Sidoli, Zuo-Fei Yuan, Kyung-Min Noh, Benjamin A. Garcia

**Affiliations:** Epigenetics Institute, Department of Biochemistry and Biophysics, Perelman School of Medicine, University of Pennsylvania, Philadelphia, PA 19104, USA; Department of Biochemistry and Molecular Biophysics, Washington University School of Medicine, St. Louis, MO 63110; Department of Biochemistry, Albert Einstein College of Medicine, Bronx, NY 10461, USA; Center for Proteomics and Metabolomics, St. Jude Children’s Research Hospital, Memphis, TN 38105, USA; European Molecular Biology Laboratory (EMBL), Genome Biology Unit, Heidelberg, Germany; Collaboration for joint PhD degree between EMBL and Heidelberg University, Faculty of Biosciences, Heidelberg, Germany

**Keywords:** histone H3.3, methylation, mouse embryonic stem cells, differentiation, H3.3K27me3

## Abstract

Histone variants, such as histone H3.3, replace canonical histones within the nucleosome to alter chromatin accessibility and gene expression. Although the biological roles of selected histone post-translational modifications (PTMs) have been extensively characterized, the potential differences in the function of a given PTM on different histone variants is almost always elusive. By applying proteomics and genomics techniques, we investigate the role of lysine 27 tri-methylation specifically on the histone variant H3.3 (H3.3K27me3) in the context of mouse embryonic stem cell pluripotency and differentiation as a model system for development. We demonstrate that while the steady state overall levels of methylation on both H3K27 and H3.3K27 decrease during differentiation, methylation dynamics studies indicate that methylation on H3.3K27 is maintained more than on H3K27. Using a custom-made antibody, we identify a unique enrichment of H3.3K27me3 at lineage-specific genes, such as olfactory receptor genes, and at binding motifs for the transcription factors FOXJ2/3. REST, a predicted FOXJ2/3 target that acts as a transcriptional repressor of terminal neuronal genes, was identified with H3.3K27me3 at its promoter region. H3.3K27A mutant cells confirmed an upregulation of FOXJ2/3 targets upon the loss of methylation at H3.3K27. Thus, while canonical H3K27me3 has been characterized to regulate the expression of transcription factors that play a general role in differentiation, our work suggests H3.3K27me3 is essential for regulating distinct terminal differentiation genes. This work highlights the importance of understanding the effects of PTMs not only on canonical histones but also on specific histone variants, as they may exhibit distinct roles.

## Introduction

Histone proteins organize DNA at the fundamental unit of chromatin, the nucleosome. The nucleosome core particle is a protein-DNA complex consisting of 147 base pairs of DNA wrapped around an octamer of histones, containing two copies each of the histones: H2A, H2B, H3, and H4^1^. Post-translational modifications (PTMs) to histone N-terminal tails can modulate gene expression by altering nucleosome interactions to open or condense chromatin, or by recruiting chromatin modifying enzymes^2^. One of the vast array of PTMs that occurs on histones is lysine methylation, which exists in three different forms: mono- (me1), di- (me2), or trimethyl (me3)^2^. Methylation on different histone residues can elicit distinct biological outcomes. For example, the trimethyl form on histone H3K4 (H3K4me3) is found at promoter regions and is associated with gene activation, whereas the trimethyl form on H3K27 (H3K27me3), is associated with repressed chromatin^3, 4^. Another form of regulation involves replacing canonical histones within the nucleosome by histone variants, which differ by a small amount of sequence variation from their canonical counterpart, to alter gene accessibility^5^. Although modification sites are conserved between variants and their canonical counterparts, it is not evident if they have identical effects.

The histone H3 variant, histone H3.3, differs from the canonical H3 histones, H3.1 and H3.2, by 5 and 4 amino acids, respectively. H3.3 is replication-independent, whereas the expression of canonical H3 histones is tied to the cell cycle^6, 7^. Previous studies noted H3.3 enrichment at promoter regions of actively transcribed genes across various cell types, identifying a critical role for H3.3 in early development^7-13^. Moreover, H3.3 is also enriched at pericentric heterochromatin and telomeres^13^. During brain development, H3.3 was established as a critical regulator of neural stem cell proliferation and differentiation. Knockout of H3.3 in murine neural stem cells *in utero* resulted in premature and abnormal neuronal differentiation^14^. H3.3-null mouse embryos displayed growth delays and did not survive past embryonic day 6.5^15^. A zebrafish model demonstrated that reduced H3.3 incorporation into chromatin resulted in disrupted neural crest development^16^. Collectively, the study of histone variants and their modifications is crucial for a better understanding of development, as both are fundamental in regulating gene function.

To further its complexity, H3.3 has been implicated in nervous system function and disease. H3.3 accumulates with age in neuronal chromatin, accounting for over 94% of the total H3 pool in aged mice^17^. Furthermore, H3.3 turnover in neurons is required for activity-dependent transcription and mediating synaptic connectivity^17^. Mutations in H3.3 have been reported in neurodevelopmental disease and various cancers^18-21^. Studies of diffuse intrinsic pontine glioma and thalamic gliomas identified a missense mutation in canonical H3 as well as H3.3, resulting in a lysine 27 to methionine substitution (H3K27M and H3.3K27M)^19-21^. H3.3K27M mutations exhibit distinct clinical features compared to the K27M mutation on canonical H3^22^. Thus, further study on the role of H3.3 in development and disease would be valuable for our understanding about the pathological mechanism of H3.3 mutations.

Trimethylation on the canonical H3K27 (H3K27me3) has been well characterized in early development. In mouse embryonic stem cells (mESC), 75% of domains that contained the repressive H3K27me3 mark also contain the activating H3K4me3 mark. These bivalent domains are located at promoter regions of transcription factors and upon differentiation resolve into a single active or repressive mark^23^. Establishment of these bivalent domains was determined to be H3.3-dependent, as H3.3 localization was required for the H3K27 methyltransferase, Polycomb repressive complex 2 (PRC2), enrichment at these loci.

In contrast to H3K27me3, the role of H3.3K27 methylation in development remains unclear. While H3K27me3 is a well-characterized repressive mark^3^, H3.3 is enriched with PTMs associated with active transcription^8^. Thus, it is unknown what function K27 methylation would play on H3.3. However, emerging evidence suggests a critical role for H3.3K27 in early development. H3.3K27R-expressing embryos showed abnormal and reduced developmental progression, as well as a defect in heterochromatin formation in the male pronuclei^24^. This effect may be due to the inability to post-translationally modify H3.3K27, as H3.3K27R embryos have reduced H3K27me3 levels^24^. In Drosophila, H3.3K27R containing mutants showed increased expression of Polycomb group target genes *Psc* and *Su(z)2* by RNA-seq, suggesting that H3.3K27R potentially leads to de-repression of Polycomb group target genes, and may be required for proper Polycomb function^25^. Notably, other studies report that H3.3K27R-expressing pluripotent mESC have a largely undisturbed transcriptome^26^.

Despite the extensive work studying the role of H3.3 in development, the field has yet to understand the role of H3.3K27 methylation for gene regulation in this context. In this work we aimed to provide an in-depth characterization of H3.3K27me3 during stem cell differentiation. To this end, we assessed histone PTM dynamics to gain insight on the dynamic regulation of genes. Mass spectrometry (MS) analysis allows for the interrogation of PTMs and protein dynamics through stable isotope labeling and can distinguish H3.3K27 from the canonical H3K27^27, 28^. Furthermore, we utilized ChIP-seq, as well as proteomic and transcriptomic analysis of H3.3K27A mutants, to elucidate the role of H3.3K27. Through our multi-omic profiling, we find that H3.3K27 exhibits unique quantitative and biological characteristics in comparison with canonical H3K27 methylation during stem cell differentiation, highlighting the importance of distinguishing PTMs on histone variants and canonical histones.

## Experimental Methods

### Cell Culture

Unless specifically indicated as CRISPR mutant H3.3K27A or H3.3A knockout mouse embryonic stem cells (mESC), all experiments used CCE-Nanog-GFP mESC that were donated by Ihor Lemischka. Cells were cultured at 37°C in a 5% CO2 atmosphere on 10 cm plates coated with 0.1% gelatin in Dulbecco’s Modified Eagle Medium with L-glutamine, high glucose (4.5g/L), and sodium pyruvate (Corning) plus 1% Glutamax, 1% Non-essential amino acids, 15% characterized fetal bovine serum (Hyclone), 0.1 mM 2-mercaptoethanol (Gibco, Thermo Fisher Scientific 21985023), and supplemented with ESGRO leukemia inhibitory factor (LIF) (Millipore) to maintain stem cell pluripotency. Upon 70% confluency, they were differentiated to embryoid bodies using 10 uM all-trans retinoic acid (Sigma) on 10 cm petri dishes in the absence of LIF. All cell lines were tested for mycoplasma. H3.3A knockout and H3.3K27A mESC lines were initially grown on feeder cells, mouse embryonic fibroblasts (MEFs), and then passaged onto feeder-free, gelatin-coated flasks. A MEF depletion step was included to reduce feeder cell contamination during mESC harvesting. Cells were harvested on day 0 (pluripotency), day 3 and day 6 of differentiation and were snap frozen and stored at -80°C until sample processing.

To assess histone methylation dynamics, mESC were cultured in ^13^CD3-methionine-containing Dulbecco’s Modification of Eagle’s Medium (DMEM) with 4.5 g/L glucose and sodium pyruvate without L-glutamine, L-methionine, and L-cysteine (Corning), containing the same additives as described above, and was supplemented with unlabeled L-glutamine (584 mg/L), L-methionine (30mg/L), and L-cysteine (62.57 mg/L). For the labeled media, the methionine supplemented was ^13^CD3-methionine (Sigma Aldrich).

To track histone turnover, mESC were cultured in DMEM without L-lysine, L-arginine (Thermo Fischer Scientific) which was supplemented with either unlabeled L-lysine (146.2 m/L) and unlabeled L-arginine (84 mg/L), or unlabeled L-lysine and ^13^C6,^15^N4-arginine (Arg10) (Silantes). An excess of proline was added to both media (600 mg/L) to prevent the conversion of heavy arginine to heavy proline during the labeling experiment^29^. Supplements were the same as above except for 15% dialyzed fetal bovine serum.

### Histone analysis

Histones were extracted, derivatized with propionic anhydride, and digested as previously described^30^. Heavy labeled synthetic histone peptide standards (Cell Signaling Technology) (Fig. S1C) were mixed together for a final concentration of 1 pmol/ul of each peptide and derivatized. Standards were analyzed by nanoLC-MS/MS using 1ul injections. Additionally, the synthetic peptide standard mix was spiked into the histone samples for a final concentration of 250 fmol/ul as an internal normalization. All samples and standards were desalted prior to nanoLC-MS/MS analysis using in-house prepared C18 stage-tips.

Histone samples were analyzed by nanoLC-MS/MS with an Easy-nanoLC coupled to an Orbitrap Fusion mass spectrometer (Thermo Fisher Scientific). The column was packed in-house using reverse-phase 75 µm ID × 17 cm Reprosil-Pur C18-AQ (3 µm; Dr. Maisch GmbH). The HPLC gradient was: 5% to 40% solvent B (A = 0.1% formic acid; B = 80% acetonitrile, 0.1% formic acid) over 47 min, from 40% to 90% solvent B in 5 min, 90% B for 8 min. The flow rate was at 300 nL/min. Data were acquired using a data-independent acquisition method, consisting of a full scan MS spectrum (m/z 300−1100) performed in the Orbitrap at 60,000 resolution with an AGC target value of 2e5, followed by 16 MS/MS windows of 50 m/z using HCD fragmentation and detection in the ion trap. HCD collision energy was set to 28, AGC target at 1e4, and maximum inject time at 50 ms. The histone samples for the H3.3K27A experiment were analyzed by nanoLC-MS/MS with a Dionex-nanoLC coupled to an QE-HF mass spectrometer (Thermo Fisher Scientific), with the same conditions as described above. The only differences were: HCD collision energy was stepped (25, 27.5, 30), AGC target at 5e5, and maximum inject time set at auto. Histone samples were resuspended in buffer A and 1 ug of total histones was injected.

Histone data was analyzed using a combination of EpiProfile 2.0^31^, Skyline^32^, and manual analysis with Xcalibur (Thermo Fisher Scientific). The peptide relative ratio was calculated by using the area under the curve (AUC) for that particular peptide over the total AUC of all possible modified forms of that peptide. Data analysis was performed using Microsoft Excel to calculate averages and standard deviations. The AUC for each heavy labeled synthetic peptide was plotted at each concentration to generate a standard curve (Fig. S1D, E) for calculating the absolute abundance of endogenous peptides.

For ^13^CD3-methionine labeled histones, the relative abundance ratio of heavy methylation incorporation was calculated by dividing the AUC for a specific heavy labeled methylated species as the by the sum of the AUCs for all labeled and unlabeled forms of that methylated species, e.g. for H3.3K27me2^**^ the ratio is: (H3.3K27me2^**^)/(H3.3K27me2^**^ + H3.3K27me2^*^ + H3.3K27me2). Arg10 labeled histones were analyzed in the same manner, where the ratio is labeled/(labeled + unlabeled).

### Antibodies

ChIP-grade antibodies were αH3.3K27me3 (custom antibody made by Proteintech), αH3.3 (Invitrogen RM190), αH3K27me3 (Active Motif 39155), αH3 (Abcam 1791), rabbit IgG (Abcam 171870) (Fig. S5). Additional antibodies used for western blotting (WB) included αH3K27me3 (Abcam 6002), αREST (EMD Millipore 07-579), αH3K4me3 (Abcam 8580), αH3K27ac (Active Motif 39133), and αH3K9me3 (Abcam 8898).

### ChIP-seq sample preparation and data analysis

Samples were prepared for ChIP-seq as previously described^33^. Briefly, 2×10^7^ mESC (day 0 pluripotent cells and day 6 embryoid bodies) were fixed in 1% formaldehyde in PBS for 5 minutes, and quenched using 125 mM glycine incubating for 5 minutes. Two replicates were conducted per ChIP. Chromatin was sonicated in the Covaris AFA Ultrasonicator (Fisher Scientific Sonic Dismembrator Model 100) for 15 minutes at 200 power (with duty set to 5 and cycles set as 200) at 4°C. All ChIPs were performed using 600ug of sonicated lysate and 5 ug of antibody. Antibodies were conjugated to 30 uL of Protein G Magnetic Beads (Dynabeads, Thermo Fisher Scientific 10-003-D), per ChIP. Eluted samples were reverse crosslinked overnight at 65°C, and DNA was purified after treatment with RNAse A (Thermo Fisher Scientific) and Proteinase K (Thermo Fisher Scientific). Purified DNA was quantified using a Qubit dsDNA kit (Thermo Fisher Scientific) and sequencing libraries were prepared using the NEBNext Ultra DNA Library Prep Kit for Illumina (New England Biolabs). Single-end sequencing (75 bp) was performed on a NextSeq 500 platform (Illumina).

FASTQ files from separate single-end sequencing runs were filtered for a Q score of at least 30 and then concatenated for alignment to the mm10 genome with STAR (v2.5.2a) with default settings except -- outFilterMultimapNmax 20 --outFilterMismatchNmax 999. Alignments were filtered for uniquely mapping reads. BigWig files, normalized to sequencing depth (RPM), were generated for display in the UCSC genome browser. MACS2 was used to call peaks from BED files with settings -g mm --no lambda --nomodel -p 1e-2 --min-length 1000 --max-gap 5000 --call-summits --cutoff-analysis. Peak sets were then filtered for pileup <= 50, fold-enrichment >= 2, and -log10 pvalue >= 3. Peaks from ChrM and ChrUn were excluded. These filtered peak sets were used for quantitative analysis of indexed bam files with DiffBind. Profile plots and heatmaps were generated with DeepTools.

### RNA-seq sample preparation and data analysis

RNA was extracted using a RNAeasy kit (Qiagen 74014) following the manufacturer’s guidelines, including DNase treatment. Libraries were prepared using NEBNext Poly(A) mRNA magnetic isolation module and NEB Ultra Directional RNA Library kit for Illumina (New England Biolabs).

FASTQ files were aligned to the mm10 genome with STAR with default settings except -- outFilterType BySJout --outFilterMultimapNmax 20 --alignSJoverhangMin 8 --alignSJDBoverhangMin 1 --outFilterMismatchNmax 999 --alignIntronMax 1000000. After filtering for uniquely mapping reads, alignments were analyzed by HTSeq (-f sam -r name -s reverse -t exon -i gene_id --nonunique none -- secondary-alignments ignore --supplementary-alignments ignore) and DESEQ2. Genes were pre-ranked by log2(day6/day0) for analysis by GSEA.

### CRISPR editing

The *H3f3b* K27A mutant was generated following the editing strategy previously described^34, 35^. Briefly, the guide RNA (sgRNA) was cloned into the pSpCas9(BB)-2A-GFP (PX458; Addgene) vector^36^. Gene editing was performed co-transfecting 2×10^6^ *H3f3a*^-/-^ mESCs with 2 μg pSpCas9(BB)-2A-GFP plasmid containing the sgRNA and 100 μM of single-stranded oligonucleotide (ssODN) repair template (180 bp, Integrated DNA Technologies Ultramers), using electroporation (Lonza Nucleofector, program CG-104). After 48h, single-cell sorting was performed to select GFP+ cells and single clones were then expanded for genotyping and freezing. Sanger sequencing confirmed successful insertion of the homozygous K27A mutation, and the chromosomal integrity of the clone was assessed by low-coverage whole-genome DNA-seq. Cells in which *H3f3a* was knocked out but *H3f3b* retained the wild-type sequence were also created to be used as a control for gene dosage (H3.3A knockout (KO) cells). sgRNA_H3f3b_g5: CGCGCTTTTCCGAGCCGCCT Repair template: CCGGCGGCCAAGTCATTGTTCCCAGCCCTCTGCCTACCTGTAGCGGTGAGGCTTCTTCACCCCGCCGGTAGAGGGCGCGCTGGCCCGAGCCGCCTTTGTGGCCAGCTGTTTGCGGGGGGCTTTCCCACCGGTGGACTTCCTAGCGGTCTGCTTGGTTCGGGCCATTTTTTTTCACCTGCGAGA

### Proteomics analysis

Proteomics samples were prepared following ProtiFi s-trap protocol^37^ and resuspended in buffer A (0.1% formic acid). 1 ug was injected and analyzed by nanoLC-MS/MS with a Dionex-nanoLC coupled to an QE-HF (Thermo Fischer Scientific). The column was packed in-house as previously mentioned. The HPLC gradient was: 5% to 35% solvent B (A = 0.1% formic acid; B = 80% acetonitrile, 0.1% formic acid) over 90 min, from 35% to 60% solvent B over 10 min, 60% B to 95% B in 5 min, and hold at 95% B for 15 min, and back down to 2% B in 1 minute. The flow rate was at 300 nL/min. Data were acquired using a data-independent acquisition method (DIA)^38^, consisting of a full scan MS spectrum (m/z 385−1015) performed in the Orbitrap at 60,000 resolution with an AGC target value of 1e6, followed by 25 overlapping MS/MS windows of 24 m/z using HCD fragmentation and detection in the orbitrap at 30,000 resolution. HCD collision energy was set to 27, AGC target at 1e6, and maximum inject time at 60 ms. A pooled sample (1.5 ug of each sample combined) was analyzed with several DIA runs to build a chromatogram library for data analysis^39^. The only difference for those methods was the full MS for each method covered a range of only 110 m/z with 25 overlapping MS/MS windows of 4 m/z each. For data analysis, .raw files were converted to .mzML using MSConvert^40^. Then a chromatogram library was built using Walnut in EncyclopeDIA, and data was searched with the library in EncyclopeDIA^39^.

### Bioinformatics and statistical analysis

Gene ontology enrichment analysis was performed with GOrilla^41, 42^. Transcription factor motif analysis was performed using Motif Analysis of Large Nucleotide Datasets (MEME-ChIP)^43^. Transcription factor signatures were assessed using Gene Set Enrichment Analysis (GSEA)^44, 45^. Network analysis was performed using STRING^46^ and visualized using CytoScape^47^ (v 3.5.1). Unless otherwise indicated, the Student’s Type II T-test was used to determine statistical significance for histone and proteomics mass spectrometry datasets between two conditions. Data visualization was performed using R packages such as ggplot2^48^. Colors and fonts of plots were altered using Adobe Illustrator.

## Data access

All raw and processed sequencing data generated in this study have been submitted to the NCBI Gene Expression Omnibus (GEO; https://www.ncbi.nlm.nih.gov/geo/) under accession number GSE169745. All raw mass spectrometry data files have been uploaded onto Chorus (https://chorusproject.org/pages/index.html) under the project number 1715.

**More details about methods can be found in Supplementary Information**.

## Results

### H3.3K27me3 Exhibits Distinct Methylation Characteristics Compared to Canonical H3K27me3

Individually, both H3K27me3 and variant H3.3 have been described as critical regulators of early development. However, the role of methylation on H3.3 compared to canonical H3 remains poorly understood, and we aimed to start elucidating this role by first quantifying the methylation on H3.3K27. To this end, mESC were grown in pluripotency and differentiated using 10uM retinoic acid (RA) to embryoid bodies (Fig. 1A), and express expected pluripotency and differentiation markers at each cell state (Fig. S1A,B). High RA concentrations, as used here, promote differentiation to the neural lineage^49, 50^. Interestingly, the mono-, di-, and tri-methylated forms of methylation on H3.3K27 are detected in pluripotency, and all decrease upon differentiation. The tri-methylated H3.3K27 (H3.3K27me3) is present at about one-tenth the level of the tri-methylated canonical H3 (H3K27me3) in pluripotency (Fig. 1B). Although both H3K27me3 and H3.3K27me3 decrease over differentiation, the degree of reduction differs, which is apparent by comparing methylation levels between the time points. The change in K27me3 over differentiation differs significantly between H3.3 and H3, for both time points (Fig. 1C).

**Figure 1.**
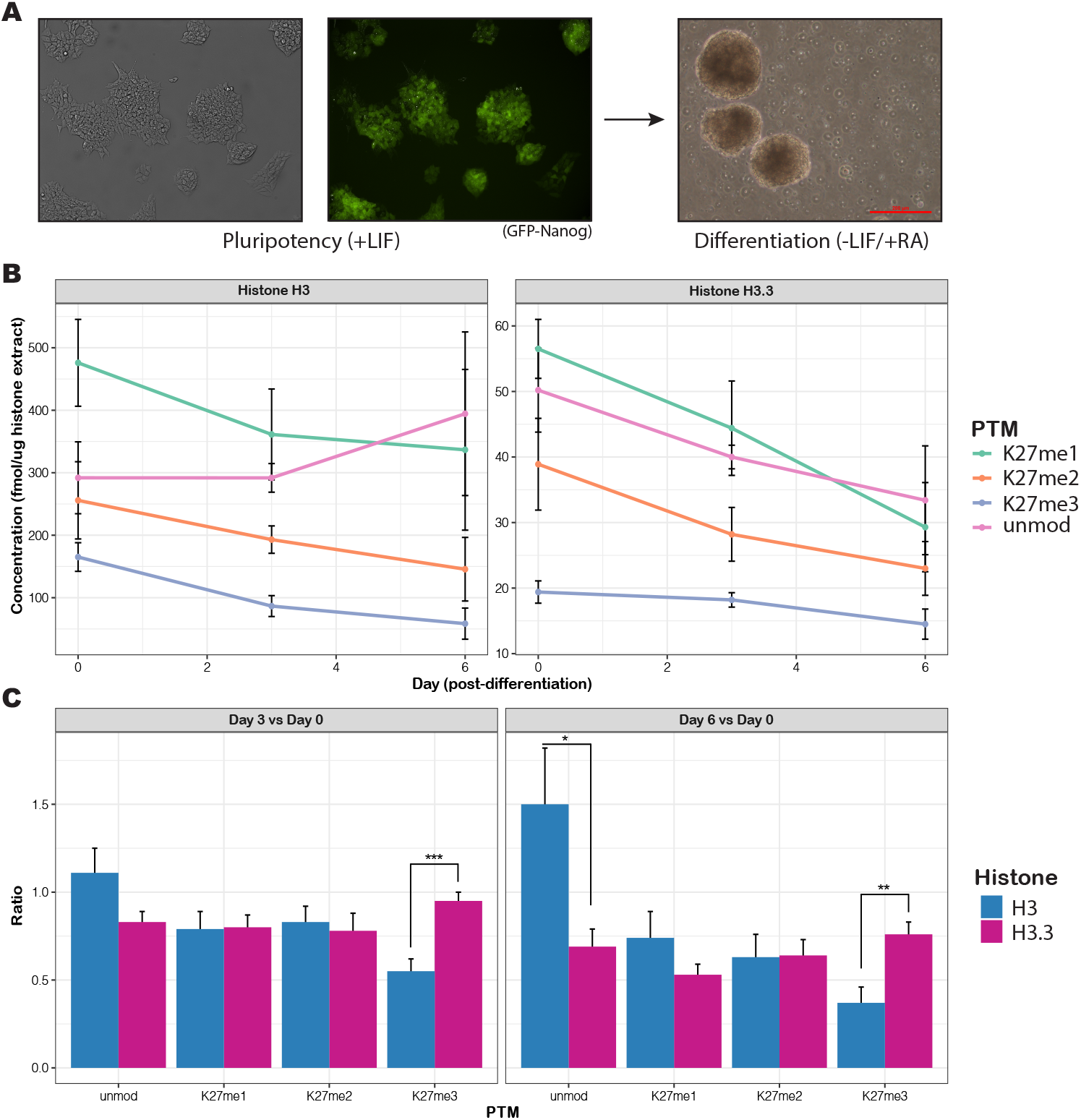
H3.3K27 and H3K27 methylation steady state levels decrease over differentiation. **A)** Mouse embryonic stem cells (mESC) were grown in pluripotency (left two images) with leukemia inhibitory factor (LIF) and express GFP-Nanog, a pluripotency marker. Cells were differentiated with 10 uM retinoic acid (RA) to form embryoid bodies (right image). **B)** Absolute quantification of methylation levels on H3.3K27 (right) and H3K27 (left). Time points are day 0 (pluripotency) and day 3 and day 6 of differentiation. The concentration was calculated from a standard curve of synthetic peptides (Fig. S1) and indicates the concentration in femtomoles per microgram of histones injected on the mass spectrometer. Biological triplicates were analyzed, and the standard deviation of the mean is indicated. **C)** The ratios of the methylation levels on day 3 versus day 0, calculated for both H3 and H3.3 (all biological replicates) is plotted on the left, and the ratios of the methylation levels on day 6 versus day 0, calculated in the same manner, is plotted on the right. A Type III T-test (heteroscedastic) was used to calculate significance. P-value < 0.05, ^*^; p-value < 0.01, ^**^; p-value < 0.001, ^***^.

Next, we applied a labeling approach to track new methylation incorporation over time. mESC were grown in ^13^CD3-methionine containing media, either in pluripotency or differentiation. MS analysis detected ^13^CD3-methyl incorporation into histones, resulting in a +4 Da isotopic mass shift per labeled methyl group. Incorporation of heavy labeled methyl groups on H3.3K27me3 was observed (Fig. S2A). Methylation incorporation ratios were plotted over time to depict the distribution of differentially labeled species in a given methylation state, both in pluripotency (Fig. S2B), and after differentiation (Fig. 2A). H3.3K27me3^***^ (number of asterisks indicate the number of heavy labeled methyls) exhibits slower heavy methylation incorporation when compared to a known active mark, H3K4me3^***^, but displays a similar trend of incorporation for repressive marks such as H3K27me3^***^ and H3K9me3^***^ (Fig. 2A). Overall, H3.3K27me3^***^heavy methylation incorporation over differentiation clusters with known repressive marks such as H3K9me3^***^, H3K27me3^***^, and H4K20me3^***^ (Fig. 2B). Interestingly, the amount of heavy labeling incorporation on H3.3K27me3^***^ is actually less at each time point over differentiation when compared to the canonical H3K27me3^***^. At the day 1 timepoint, there is about 40% heavy label incorporation for H3.3K27me3^***^, but about 50% for H3K27me3^***^ (Fig. 2A).

**Figure 2.**
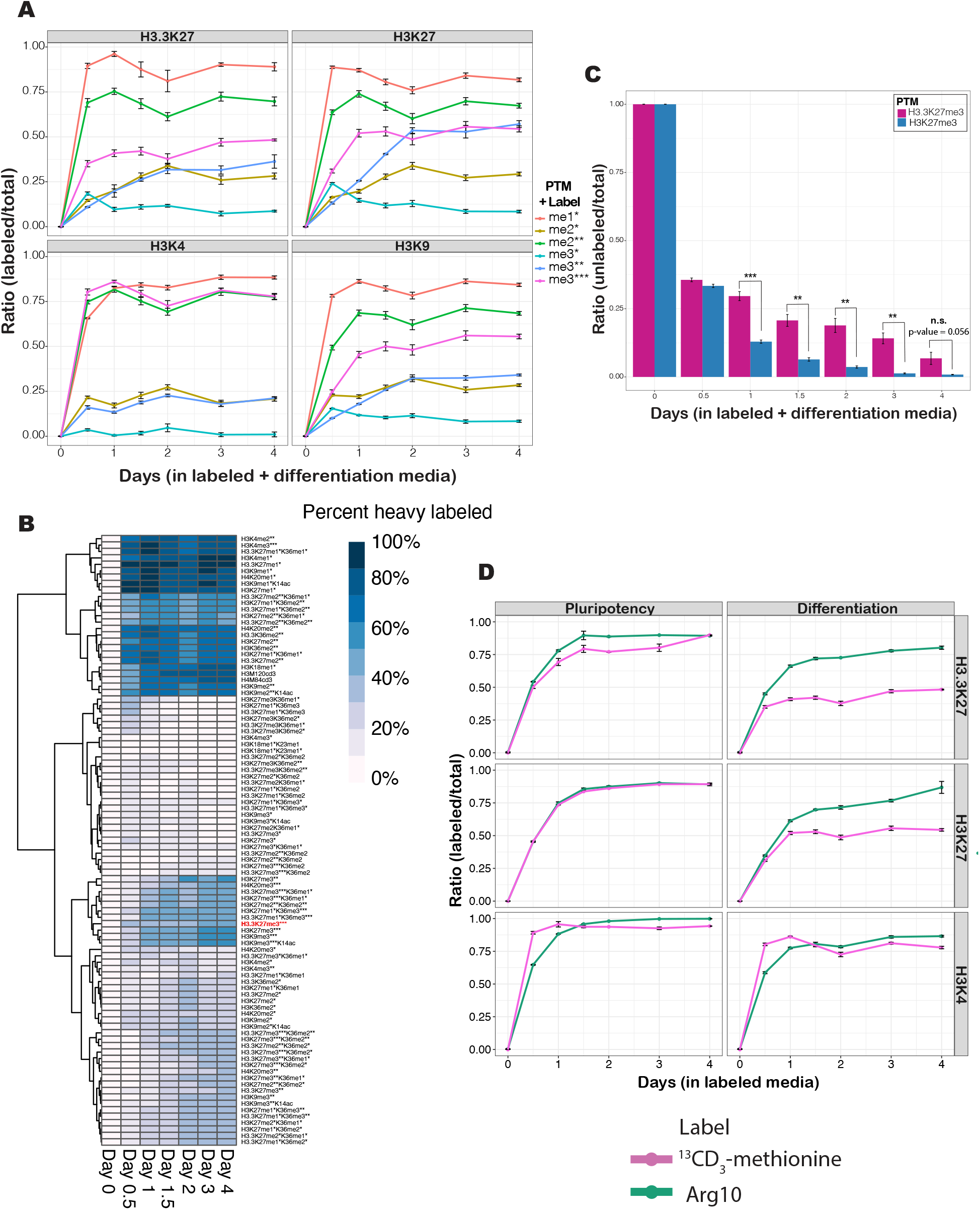
H3.3K27me3 is maintained compared to H3K27me3 over differentiation, and is independent of histone turnover. **A)** Heavy methylation incorporation (^13^CD3-methionine) is plotted over time points in labeling/differentiation media for the histone marks H3.3K27 (top left), H3K27 (top right), H3K4 (bottom left) and H3K9 (bottom right). Time points collected were at 0 (unlabeled control), 0.5, 1, 1.5, 2, 3, and 4 days after switching to labeled media. The ratio of heavy labeling is a specific heavy labeled methylated species over the sum of all labeled and unlabeled forms of that methylated species. Asterisks indicate the number of methyl groups that are heavy labeled. The mean of biological triplicates is plotted with the standard deviation of the mean. **B)** Heavy methylation incorporation is plotted over differentiation and labeling time points (^13^CD3-methionine), with PTMs clustering based on their incorporation trends. Light purple indicates 0% heavy labeled, and dark blue indicates 100% heavy labeled. H3.3K27me3^***^ is bolded and highlighted in red. **C)** The mean ratio of unlabeled peptide is plotted over the labeling and differentiation time points, with the standard deviation of the mean indicated. A Student’s Type II T-test was used to calculate significance. P-value < 0.05, ^*^; p-value < 0.01, ^**^; p-value < 0.001, ^***^. **D)** The ratio of labeling for either methylation (^13^CD3-methionine labeling, indicated in pink) or histone turnover (^13^C6,^15^N4-arginine labeling, labeled as “Arg10” and indicated in green) is plotted over time points in labeling media. The ratio for methylation labeling is calculated for the fully labeled tri-methyl species (me3^***^), calculated the same as for panel A. The arginine labeling ratio is the labeled peptide over the sum of the labeled and unlabeled peptide. The heavy labeling ratios are plotted separately for H3.3K27, H3K27, and H3K4, in both pluripotency and differentiation. Biological triplicates were analyzed, with the standard deviation of the mean indicated.

Dynamic methylation can be tracked through the disappearance of the unlabeled methyl groups as they are replaced by labeled ones over time. Surprisingly, unlabeled H3.3K27me3 is higher than unlabeled H3K27me3 at each time point, indicating that methylation is being maintained on H3.3K27 over these early differentiation time points (Fig. 2C). The hybrid labeled marks identified in this experiment may offer unique insights about chromatin that is changing. For example, the H3.3K27me3^**^ was possibly H3.3K27me1 during pluripotency, and over the differentiation process in labeled media, it acquired two heavy-labeled methyls, turning it to H3.3K27me3^**^ (Fig. 2A). On H3, the amount of K27me1 that changed to K27me3 (H3K27me3^**^) is equal to amount of newly deposited H3K27me3^***^ by day 2 of differentiation (Fig. 2A). However, this is not the case for H3.3, as the H3.3K27me3^**^ failed to equate to H3.3K27me3^***^ incorporation over this differentiation time course (Fig. 2A). Despite exemplifying decreased incorporation of new heavy-labeled methylation over differentiation, H3.3K27me3 is not observed to be enriched in the most heterochromatic fraction by salt fractionation (Fig. S3). Overall, there are quantitative differences between methylation on the variant H3.3K27 compared to on canonical H3K27.

### Histone Methylation Dynamics are Independent of S-Adenosyl Methionine Labeling and Histone Turnover

Differences in heavy methylation incorporation between pluripotency and differentiation could be due to the changes in the labeling level of s-adenosyl methionine (SAM), the methyl-donating substrate of histone methyltransferases. While SAM becomes rapidly labeled to levels over 90% within 1-3 hours in ^13^CD3-methionine media for both cell states, labeling plateaus at about 95% by 24 hours in pluripotent mESC (Fig. S4A). However, in differentiating cells a decrease in labeled SAM is observed after 24 hours of differentiation, which may be due to a different upstream metabolite contributing to SAM^51^. Indeed, a 15-fold increase in methyl-tetrahydrofolate (mTHF) in day 3 differentiated embryoid bodies as compared to pluripotent mESC has been reported^52^. Thus, our observed decrease in labeled SAM in differentiation may be due to an increase in the unlabeled mTHF precursor pool and may account for the dip in methylation incorporation observed around day 2 (Fig. 2A). The difference in SAM labeling between pluripotency/differentiation was used to normalize the methylation incorporation ratios for H3.3K27. However, heavy methylation incorporation remains faster in pluripotency compared to differentiation (Fig. S4B), consistent with the fact that chromatin in pluripotent mESC is more open and accessible to labeling^53^.

Histone turnover is thought to mediate dynamics of histone PTMs^54^. To monitor histone turnover, mESC were grown in ^13^C6,^15^N4-arginine (Arg10). Time points were collected to track heavy arginine incorporation into newly synthesized histones, resulting in a +10 Da mass shift. Similar turnover was observed for both H3.3K27me3 and H3K27me3, which was slower than the turnover of the active mark H3K4me3 (Fig. S4C). Histones modified with active marks have been reported to exhibit fast turnover^28^, potentially because those histones are at more accessible genomic loci with different nucleosome replacement rates^55^. Histone turnover is faster in pluripotency compared to differentiation (Fig. S4C), since the differentiating cells are dividing less at these timepoints and will ultimately stop dividing. In pluripotency, methylation incorporation trends very similarly with histone turnover. For the active mark H3K4me3, even in differentiating cells the methylation incorporation is consistent with the turnover of the histone containing this mark. However, for the repressive mark H3K27me3 and the variant H3.3K27me3, methylation incorporation is independent from histone turnover in differentiation (Fig. 2D).

### H3.3K27me3 Exhibits Distinct Genomic Localization and Corresponding Gene Expression Profiles Compared to H3K27me3

Having observed differences in the quantitative characteristics of methylation on variant H3.3K27 and canonical H3K27 during mESC differentiation, we next sought identify any differences in their genomic localization and association with gene expression. We generated custom antibodies against H3.3K27me3 for ChIP-seq. Due to the 92% sequence similarity between the H3.3K27 and H3K27, various quality control experiments were necessary to validate their specificity for H3.3K27me3 over H3K27me3. Dot blots against the target H3.3K27me3 peptide and possible off-target peptides identified a purified IgG H3.3K27me3-specific antibody (Fig. S5A). A peptide competition assay provided further evidence of antibody specificity based on decreased signal intensity (Fig. S5B). Mono-nucleosome immunoprecipitation as well as ChIP indicate that the custom αH3.3K27me3 antibody enriches for H3.3K27me3 (Fig. S5C, D). Moreover, MS analysis confirmed 7-fold increased recognition of H3.3K27me3 with αH3.3K27me3 antibody, when compared to canonical H3K27me3 (Fig. S5E). The commercial H3K27me3 antibody (Active Motif), recognizes the canonical H3K27me3 about 5 times better than the H3.3K27me3 (Fig. S5F, G).

To localize H3.3K27me3 in the genome and clarify its effect on gene expression, RNA-seq and ChIP-seq using the αH3.3K27me3 antibody were performed at two time points: day 0 mESC (pluripotency) and day 6 post-treatment with RA (differentiation). ChIP-seq for the canonical H3K27me3, H3.3, and H3 were performed in parallel. RNA-seq biological replicates cluster together by PCA, but time points are separated, and differential gene expression is observed between cell states (Fig. S6). The ChIP-seq PCA similarly shows clustering of biological replicates but separation of ChIP targets and timepoints (Fig. S7A).

To investigate the relationship between changes in gene expression and histone methylation, we stratified genes based on differential expression and then compared ChIP-seq signals across the genes in each group. In pluripotent cells, H3K27me3 shows enrichment around the transcription start site (TSS) of genes destined for upregulation at day 6 of differentiation (Fig. 3A), consistent with previous literature characterizing H3K27me3 as a repressive mark that keeps differentiation genes silenced in pluripotent cells^3^. Interestingly, H3.3K27me3 shows only modest enrichment at those genes, and not to the same degree as canonical H3K27me3 (Fig. 3A). Next, ChIP-seq signals were compared across genes categorized by their relative expression level (top quintile, middle 3 quintiles, and bottom quintile) within the different cell states. As expected, H3K27me3 was enriched around the TSS of genes with low and medium levels of expression but depleted from those with high expression (Fig. 3B). In contrast, H3.3K27me3 did not show a strong correlation with gene expression levels, except for perhaps a minor enrichment across the bodies of highly expressed genes in differentiated cells (Fig. 3B). This provided clear evidence that gene repression likely relies more on canonical H3K27me3. Gene activation, on the other hand, has been associated with deposition of H3.3^8^. Accordingly, the ChIP-seq profile for total H3.3, but not canonical H3, demonstrated a striking positive correlation with gene expression level (Fig. S7B). Finally, we analyzed ChIP-seq signals at CpG islands, which have regulatory significance and have been shown to recruit PRC2^56^. We noted the presence of H3K27me3 but not H3.3K27me3 at these genomic regions (Fig. S7C). To correlate gene expression with histone marks, we plotted the fold change of the ChIP-seq peak signal (day 6/day 0), specifically at the promoter regions, against the fold change of the RNA-seq expression data. Genes marked with higher levels of H3K27me3 on day 6 of differentiation exhibit decreased expression levels (Fig. 3C, left). The histone variant H3.3 shows the opposite correlation, as genes marked with higher levels of H3.3 on day 6 of differentiation exhibit higher expression levels (Fig. 3C, right). Interestingly, changes in K27me3 on H3.3 exhibit neither the same strong positive correlation with changes in gene expression as H3.3 nor the same strong negative correlation as H3K27me3 (Fig. 3C, middle). There is a minor negative correlation with gene expression for H3.3K27me3; however, there is also a small population of highly expressed genes that are marked with low amounts of H3.3K27me3.

**Figure 3.**
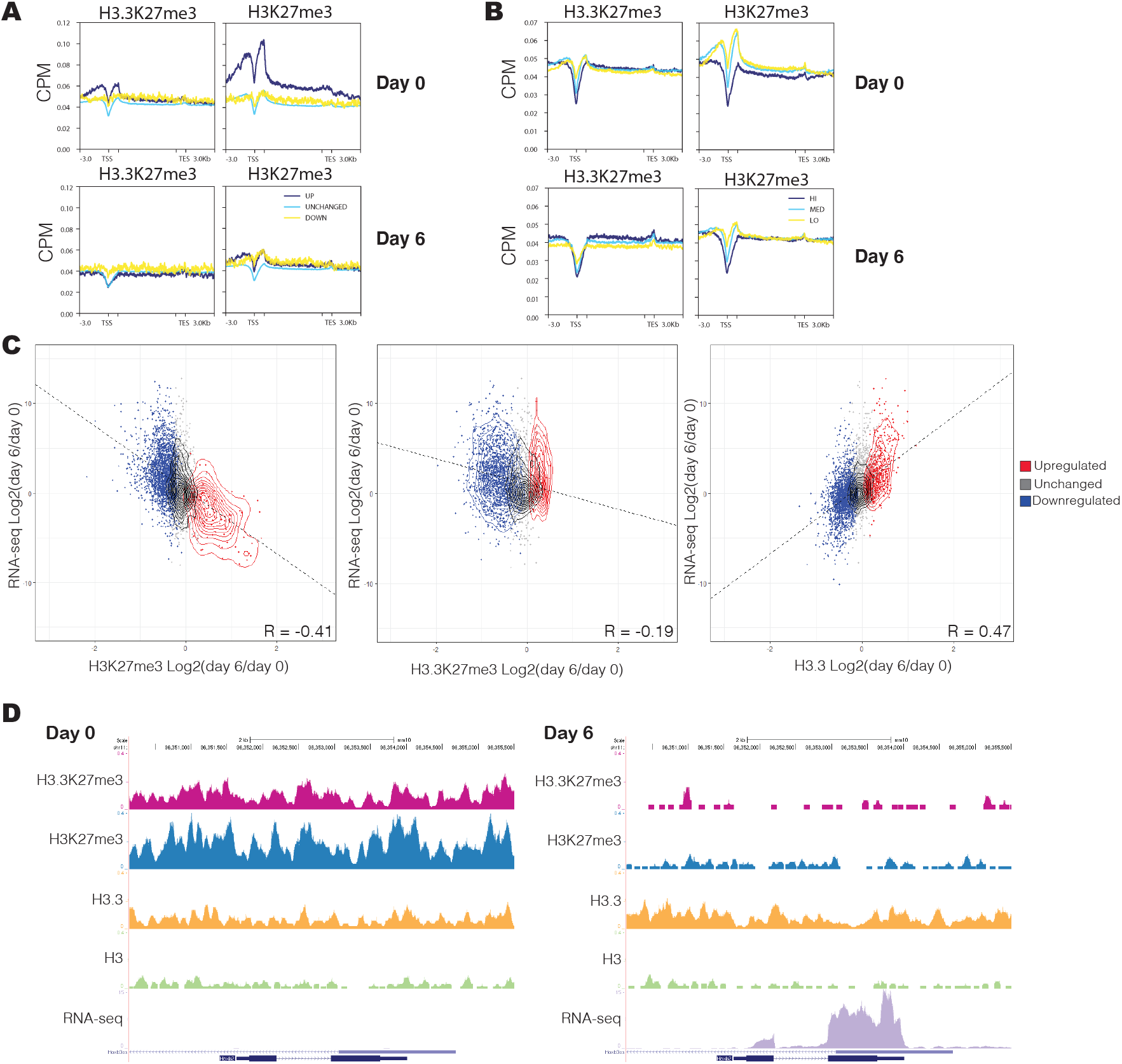
Gene expression is more negatively correlated with canonical H3K27me3, not H3.3K27me3. **A)** Based on RNA-seq data, genes were classified as upregulated (dark blue), downregulated (yellow), or unchanged (light blue) comparing differentiated cells (day 6) to pluripotent cells (day 0). The mean ChIP-seq signal for variant H3.3K27me3 or canonical H3K27me3 is plotted across these genes in 10 bp bins. The regions 3 kb upstream of the transcription start site (TSS) and 3 kb downstream of the transcription end site (TES) are also included. The inner tick marks demarcate 1 kb beyond the TSS and 1 kb before the TES. Inside this region, scaling has been applied to represent genes as a uniform length. For all the ChIP-seq and RNA-seq data shown here and following, two biological replicates were analyzed. The y-axis is CPM = counts per million mapped reads. **B)** Genes were stratified into quintiles based on their relative expression levels. The mean ChIP-seq signal is plotted across genes in the top quintile with the highest expression (HI, dark blue), the bottom quintile with the lowest expression (LO, yellow), and the middle three quintiles (MED, light blue). **C)** The relative change (Log2[day6/day0]) in RNA levels and ChIP peak intensity (variant H3.3K27me3, canonical H3K27me3, or total H3.3) is presented as a scatterplot. Each point represents a gene. ChIP-seq peaks were required to occur within 10 kb of the TSS to be associated with a gene and its RNA measurement. In cases where a single gene was associated with multiple ChIP peaks, the peak closest to the TSS (and then the most intense peak in the case of ties) was selected. Points are color coded by the trend in the ChIP-seq data (upregulated peaks in red, downregulated peaks in blue, and unchanged peaks in gray, based on FDR < 0.05 and fold-change > 0 or < 0). Contour traces are also plotted for clarity. The Pearson correlation coefficient is indicated. **D)** ChIP-seq and RNA-seq tracks for H3.3K27me3 (pink), H3K27me3 (blue), H3.3 (yellow), and H3 (green) are visualized across the Hoxb2 gene using UCSC Genome Browser^98^ (http://genome.ucsc.edu/). The RNA-seq track is shown as well. Tracks are shown for both pluripotency (day 0) and differentiation (day 6).

The role of the repressive H3K27me3 at bivalent domains has been characterized in mESCs^23^. However, it is unclear if H3.3K27me3 is found at these bivalent domains as well. We observe that for genes that have been previously characterized as being bivalent in mESC^57^, such as Hoxb2^23^, high levels of H3K27me3 exist at their TSS in pluripotent mESC. These genes are marked with H3.3K27me3 as well, although to a lesser extent (Fig. S7D, Fig. 3D). Upon differentiation, the methylation on both H3K27 and H3.3K27 is removed (Fig. S7D), and the gene transitions from repressed to transcribed (Fig. 3D).

### H3.3K27me3 is Enriched at Lineage Specific Genes and FOXJ2/3 Transcription Factor Binding Motifs

Next, we assessed the distribution of H3.3K27me3 across the genome. H3.3K27me3 peaks in pluripotency and differentiation were enriched for promoters and distal intergenic regions, respectively (Fig. 4A). H3K27me3 peaks, regardless of cell state, generally occurred at promoter regions (Fig. 4A). While ChIP-seq peaks at promoter regions showed both up- and downregulation of H3K27me3 at day 6, there was a bias for the downregulation of H3.3K27me3 peaks at promoters with an upregulation at distal intergenic regions (Fig. S8A). However, this was not a result of interplay with enhancers, as H3.3K27me3 peaks at distal intergenic regions did not overlap with previously published^58^ enhancer marks (Fig. S8B).

**Figure 4.**
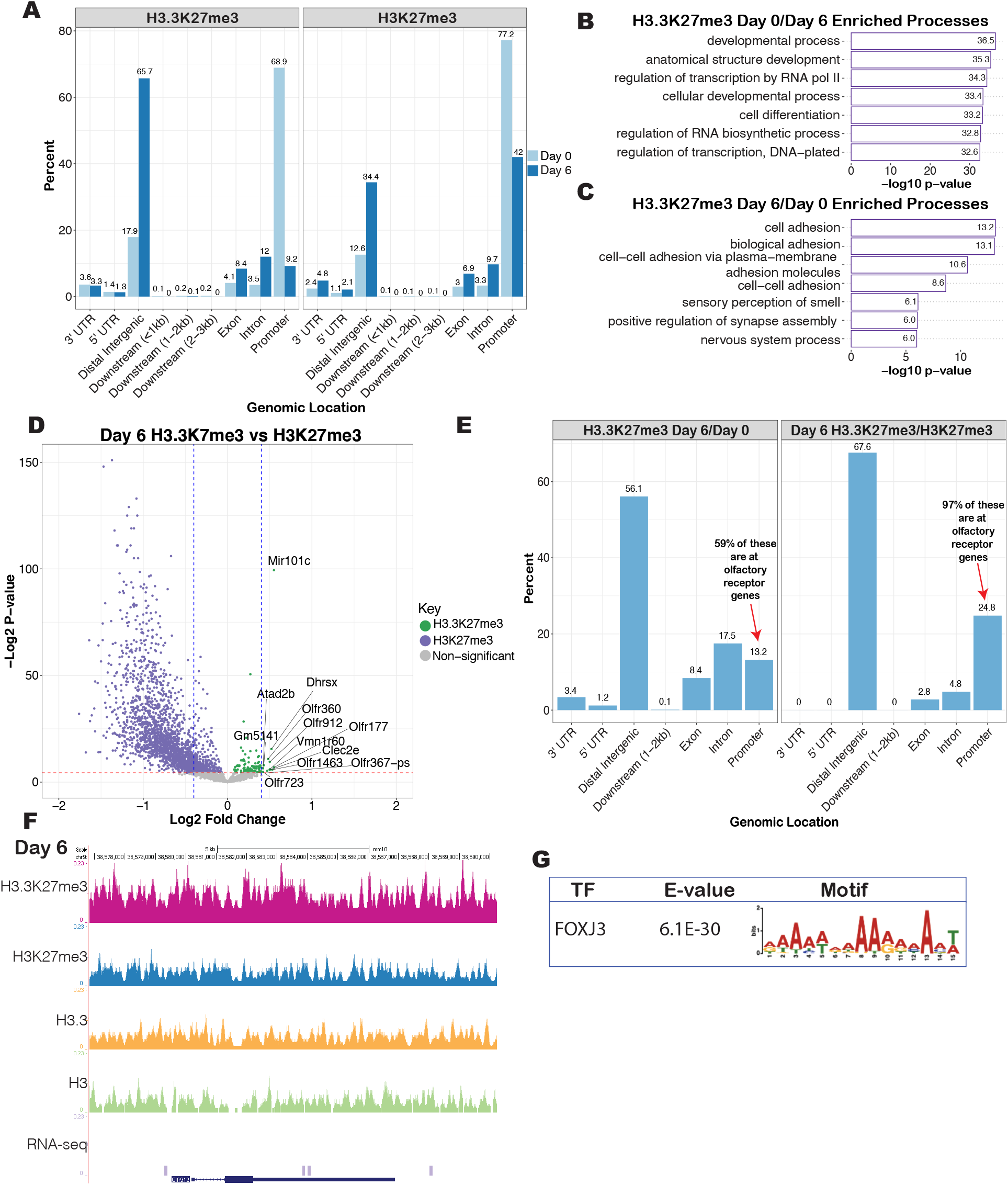
H3.3K27me3 is found at promoters and distal intergenic regions, and is enriched at lineage specific genes during differentiation. **A)** Genomic distribution of ChIP-seq peaks that are more intense on day 0 compared to day 6 (light blue) or more intense on day 6 compared to day 0 (dark blue) for H3.3K27me3 (left plot) and H3K27me3 (right plot). The counts are graphed as a percent of the total number of peaks detected for that ChIP. **B)** Gene ontology enriched processes for H3.3K27me3 ChIP-seq peaks that are more intense on day 0 (in mESC pluripotency) compared to day 6 of differentiation. Gene ontology analysis was performed using GOrilla ^41, 42^. **C)** Gene ontology enriched processes for H3.3K27me3 ChIP-seq peaks that are more intense on day 6 of differentiation compared to day 0, also analyzed with GOrilla. **D)** Volcano plot of the ChIP-seq data for day 6 of differentiation plotting the log2 fold change H3.3K27me3/H3K27me3 on the x-axis and the -log2 t-test p-value on the y-axis, for promoter region peaks. The red dotted line indicates a p-value of 0.05 (-log2 p-value = 4.32). The blue dotted lines indicate a log2 fold change of -0.4 and +0.4. Green dots indicate peaks significantly enriched with H3.3K27me3, while purple dots indicate peaks significantly enriched with H3K27me3. The volcano plot is asymmetric because there are more regions highly marked in H3K27me3 than H3.3K27me3. **E)** The genomic localization counts for the genes from panel C were plotted, stratified by comparison between time point or between histone PTMs. The counts are plotted as a percent of the total peaks detected. A total of 770 genes were identified in the gene ontology that had increased H3.3K27me3 at day 6 compared to day 0, 102 of which were at promoter regions specifically, and 60 of those promoter regions were at olfactory genes ((60/102)^*^100 = 58.8%). A total of 145 genes were identified in the gene ontology that had increased H3.3K27me3 compared to H3K27me3 at day 6, 36 of which were at promoter regions specifically, and 35 of those promoter regions were at olfactory genes ((35/36)^*^100 = 97.2%). **F)** The peak tracks for H3.3K27me3 (pink), H3K27me3 (blue), H3.3 (yellow), and H3 (green) ChIP-seq across the Olfr912 gene, as a representative olfactory receptor gene, as well as the RNA-seq expression levels, is visualized using UCSC Genome Browser ^98^ (http://genome.ucsc.edu/). **G)** The differential H3.3K27me3 peaks that were enriched at day 6 of differentiation compared to day 0 (pluripotency) were analyzed by MEME-ChIP ^43^ and the motif identified associated with FOXJ3 binding is shown. The E-value is the “enrichment value” (significance value for MEME-ChIP).

Genes with increased H3.3K27me3 peaks in pluripotency are enriched for processes involving development, regulation of transcription, and differentiation (Fig. 4B). H3K27me3-marked genes showed similar process enrichment in pluripotency (Fig. S8C). While H3K27me3 remains enriched at general development and differentiation genes at day 6 of differentiation (Fig S8D), H3.3K27me3-marked genes are enriched for lineage-specific processes, such as adhesion, sensory perception of smell, regulation of synapse assembly, and nervous system process (Fig. 4C). Moreover, H3.3K27me3 is highly enriched for olfactory receptors in both cell states (Fig. S8E-F, Fig. 4D), where 59% of the promoter proximal H3.3K27me3 peaks with higher abundance in differentiated cells occur at olfactory receptor genes (Fig. 4E). These olfactory genes comprise 97% of the promoter regions more highly marked by H3.3K27me3 compared to H3K27me3 at day 6 (Fig. 4E), and have no detectable expression (Fig. 4F).

To further explore potential regulatory effects of H3.3K27me3, motif analysis was conducted using Motif Analysis of Large Nucleotide Datasets (MEME-ChIP)^43^. H3.3K27me3 peaks having higher abundance at day 6 of differentiation were enriched for the FOXJ3 transcription factor binding motif (Fig. 4G). MEME-ChIP analysis for H3K27me3 differential peaks did not show this enrichment (Fig. S9A).

### Proteomic and Histone PTM Dysregulation in H3.3K27A mESC

Given the identified quantitative and biological distinctions between H3.3K27me3 and H3K27me3, we sought to test whether differentiation would be disrupted if methylation on H3.3K27 was lost. To this end, we generated H3.3K27A mESCs that substitute lysine with a non-modifiable alanine (*H3f3a* knocked out but *H3f3b* is K27A) (Fig. S10) using our previously established platform^35^. The control cell line used for comparison have *H3f3a* knocked out but *H3f3b* is wild-type (H3.3A KO cells), to specifically focus on the effect of the lysine 27 residue mutation. Obvious phenotypic differences were not observed during pluripotency or differentiation in H3.3K27A and control cell lines, however, alterations in pluripotency and differentiation markers were detected (Fig. S11).

Proteomic analysis of H3.3K27A mESC revealed significant dysregulation, particularly at day 6 of differentiation (Fig. S12A, Fig. 5A). Gene ontology (GO) analysis of upregulated proteins in H3.3K27A cells during day 6 revealed an enrichment for cell adhesion, in alignment with the GO enrichment of H3.3K27me3 at day 6 as found in our ChIP-seq data. Additionally, the H3.3K27A differentiating cells show an upregulation of proteins involved in development, anatomical structure morphogenesis, and glial cell differentiation (Fig. 5B). Downregulated proteins in the H3.3K27A cells include those involved with protein and vesicle transport (Fig. 5C).

**Figure 5.**
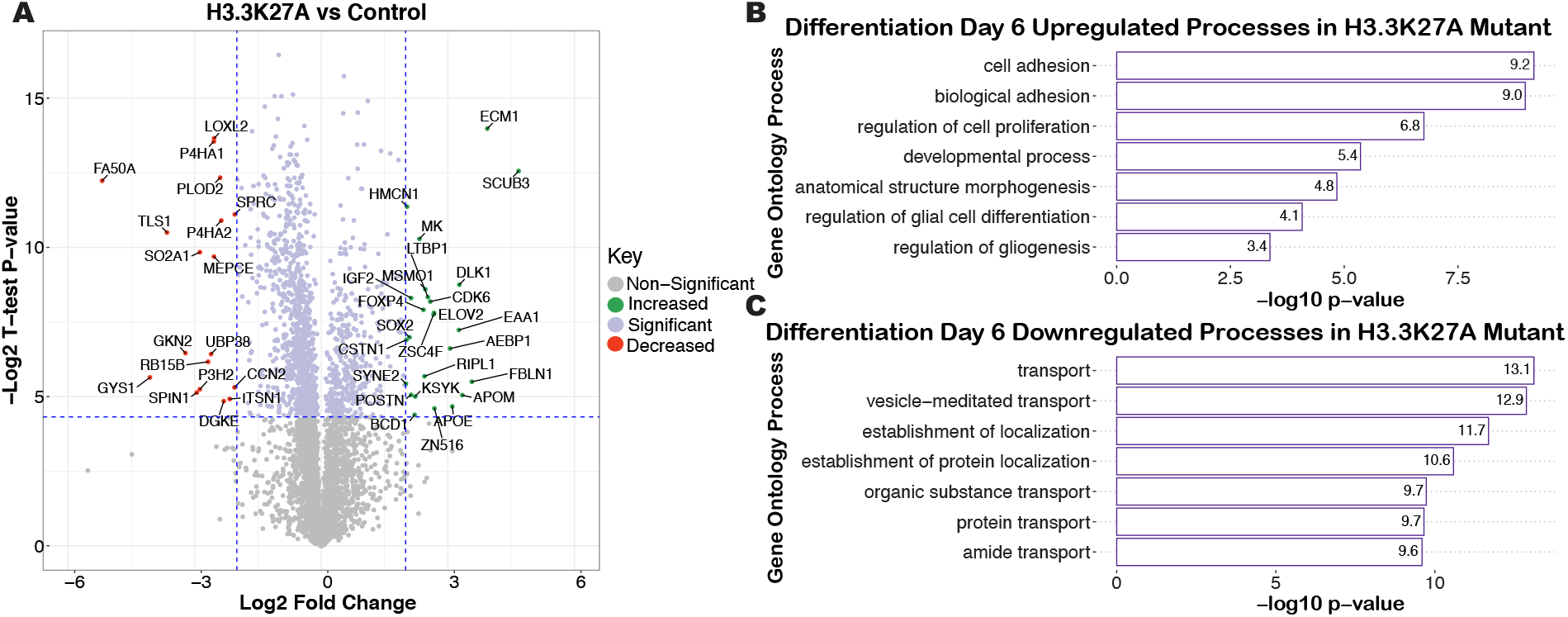
Loss of methylation at H3.3K27 results in proteomic dysregulation during differentiation. **A)** Volcano plot of proteomics data plotting the log2 fold change of H3.3K27A/control cells on day 6 of differentiation, and the -log2 t-test p-value is plotted on the y axis. The horizontal blue line indicates a - log2 p-value of 4.32 (equivalent to a p-value of 0.05), so anything above that line is statistically significant. The vertical blue lines indicate a log2 fold change of +/-2. Points indicated in green were significantly increased in the H3.3K27A mutant cells, above a log2 fold change of 2. Points indicated in red were significantly decreased in the H3.3K27A mutant (and thus increased in the control), below a log2 fold change of -2. Indicated in purple are all the other statistically significant proteins, regardless of a fold change cutoff. **B)** Gene ontology analysis was performed using GOrilla ^41, 42^ on the proteins increased in the H3.3K27A cells compared to the controls on day 6 of differentiation. **C)** Gene ontology analysis was performed using GOrilla ^41, 42^ on the proteins decreased in the H3.3K27A cells compared to the controls on day 6 of differentiation.

Proteomic changes in H3.3K27A cells may be linked to alterations in epigenetic regulation due to the direct loss of methylation or acetylation at K27, or via indirect changes in histone modifications at other residues due to an interplay with modifications at H3.3K27. To interrogate whether proteins with dysregulated expression in the H3.3K27A cells would have been marked with H3.3K27me3, we plotted the H3.3K27me3 ChIP-seq signal across the TSSs of these proteins in differentiation. However, these proteins were not highly marked with H3.3K27me3 at their TSS, suggesting the change in protein expression may not be directly due to the loss of methylation on H3.3K27 (Fig. S12B). Additionally, genes marked by H3.3K27me3 in differentiation, as determined by ChIP-seq, showed variable protein expression trends upon the loss of methylation in the H3.3K27A cells (Fig. S12C).

Decades of research have characterized crosstalk between histone modifications and its significance in gene regulation^59^. Thus, we hypothesized that this proteomic dysregulation may be due to a disruption in the balance between H3.3K27me3 and other histone PTMs that impact gene expression. Through MS analysis, we confirmed the presence of the K27A mutation based on detection of the unmodified H3.3K27A mutant peptide without detection of the K27me3 or K27ac modifications (Fig. S13A). The H3.3K27A cells exhibited altered histone H3 PTMs across both pluripotency and differentiation, with increased changes observed at day 6 of differentiation (Fig. 6A). In pluripotency, the H3.3K27A mESCs exhibit a striking depletion in H3K4me2/3 (Fig. S13B). In differentiation there is a significant increase in active marks, such as H3K4me3 and H3K27ac, and a depletion in the repressive H3K27me2/3 in the H3.3K27A cells (Fig. 6B).

**Figure 6.**
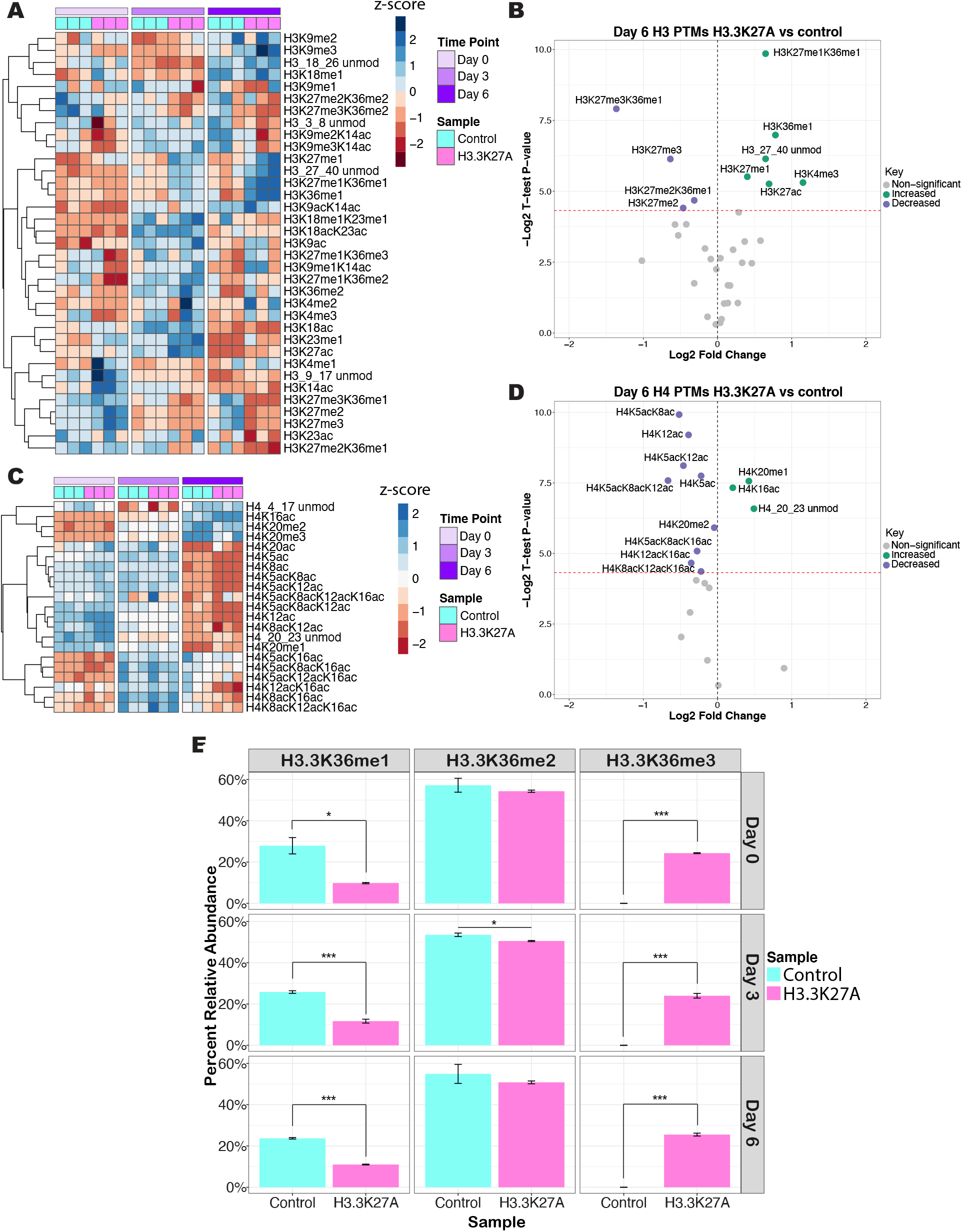
Loss of methylation at H3.3K27 results in global alterations of histone post-translational modifications. **A)** The z-scores of the relative abundances of the detected histone H3 PTMs are plotted with hierarchal clustering. The heatmap is separated by time point: day 0 (pluripotency, indicated in light purple), day 3 of differentiation (purple), and day 6 of differentiation (dark purple). The three biological replicates for the H3.3K27A mutant cells are indicated in pink and the three biological replicates for the control cells are indicated in turquoise. **B)** A volcano plot of the H3 PTMs on day 6 of differentiation. The log2 fold change of H3.3K27A/control is plotted with the -log2 t-test p-value on the y-axis. The red dotted line indicates a -log2 p-value of 4.32 (equivalent to a p-value of 0.05). Green points are significantly increased in the H3.3K27A cells, and purple points are significantly decreased in the H3.3K27A cells. **C)** The z-scores of the relative abundances of the detected histone H4 PTMs are plotted with hierarchal clustering. The heatmap is organized in the same way as described for panel A. **D)** A volcano plot of the H4 PTMs on day 6 of differentiation, organized as in panel B. **E)** The relative abundance of H3.3K36me1/me2/me3 in the H3.3K27A cells and control cells is plotted for the three time points, day 0 (pluripotency) and day 3 and 6 of differentiation. To allow for a direct comparison between H3.3K27A and control cells, in the control cells the total K36 methylation was calculated, including any peptides that also carry K27 methylation. The mean of biological triplicates is plotted, with the standard deviation of the mean. A Student’s Type II T-test was used to calculate significance. P-value < 0.05, ^*^; p-value < 0.01, ^**^; p-value < 0.001, ^***^.

The H3.3K27A mutant cells exhibit significant alterations in histone H4 acetylation at day 6 of differentiation (Fig. 6C), with only slight changes at day 0 (Fig. S13C). Strikingly, there is a global depletion in H4 acetylation at day 6, with the one exception of H4K16ac being increased (Fig. 6D). With the loss of methylation at K27, we observe altered K36 methylation on the H3.3K27A mutant itself across all time points. Namely, a decrease in H3.3K36me1 and a sizable increase in H3.3K36me3, with roughly 30% of the mutant containing K36me3 (Fig. 6E). This interplay between K27 and K36 methylation is consistent with previous work that indicates H3K27me3 and H3K36me3 are antagonistic^60-62^.

### Lack of Methylation on H3.3K27 Results in FOXJ2/3 Transcription Factor Target Upregulation during Differentiation, such as NRSF/REST, a Transcriptional Repressor of Terminal Neuronal Genes

Further analysis of the proteomics data by Gene Set Enrichment Analysis (GSEA)^44, 45^ revealed significant enrichment of the targets of FOXJ2 (normalized enrichment score: 2.09, p-value = 0.002), a close homolog of FOXJ3^63^, in the proteome of the differentiated H3.3K27A mutants (Fig. 7A). While the MEME-ChIP database includes motifs for FOXJ2/3, GSEA only contains a database for FOXJ2. FOXJ2/3 have similar predicted binding motifs (Fig. S9B), and extensive overlap in chromatin binding motifs for the forkhead box (FOX) transcription factor family has been previously identified^64^. While we detect enrichment of FOXJ3 binding motifs and FOXJ2 targets, we consider these enrichments to be overlapping and related. Genes enriched with FOXJ2 signature displayed increased protein expression in the H3.3K27A at day 6 of differentiation which may be due to increased access of FOXJ2 to these binding sites (Fig. 7A). As FOXJ2/3 have been characterized to be transcriptional activators^65 66^, it is consistent that increased chromatin accessibility leads to increased expression of targets. The upregulated FOXJ2 targets are proteins important in cell cycle control and transcriptional regulation (Fig. 7B). This enrichment of FOXJ2 targets was confirmed on the transcript level by GSEA of RNA-seq data from the H3.3K27A cells (Fig. 7C).

**Figure 7.**
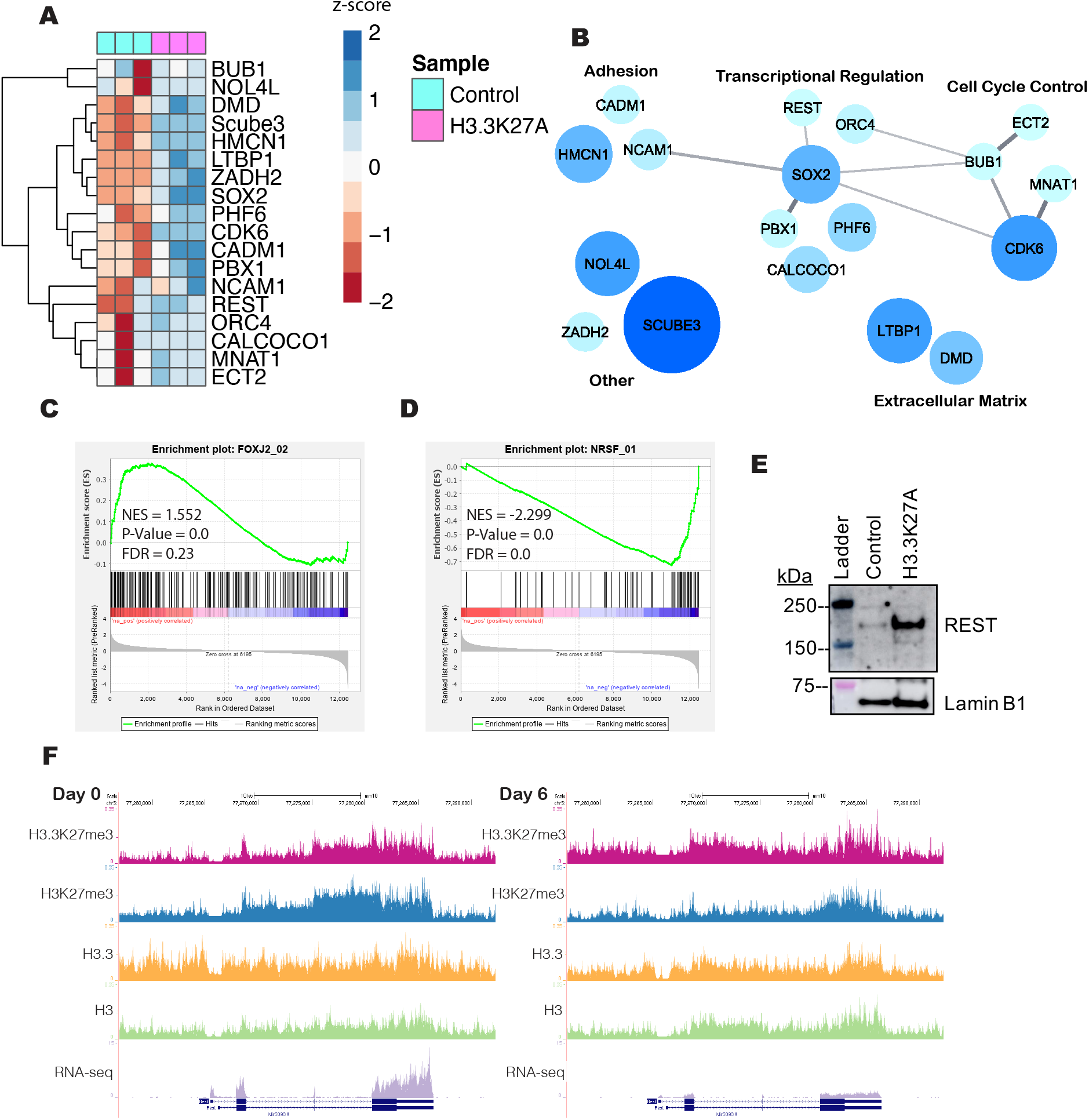
H3.3K27me3 during differentiation may regulate FOXJ2/3 binding to target genes such as REST. **A)** The upregulated targets of FOXJ2 on day 6 of differentiation as identified by GSEA analysis ^44, 45^ are visualized as a heatmap of their z-scores with hierarchal clustering. The H3.3K27A samples are indicated in pink, and the control cells are indicated in turquoise, in biological triplicate. A total of 54 FOXJ2 targets were detected in the proteomics analysis, but this subset contributed the most to the significance of the FOXJ2 transcription factor signature enrichment by GSEA. **B)** The targets in panel A are visualized as a network diagram. Bubble size and color are indicative of protein enrichment in the H3.3K27A cells compared to control cells (bigger the bubble, darker the blue means higher fold change enrichment). Proteins with known interactions are connected via a gray line, with line thickness indicative of protein interaction score as determined by STRING ^46^ (thicker line indicates higher interaction score). Proteins are grouped by functional annotation. **C)** The differentially expressed genes between H3.3K27A cells and control cells as determined by RNA-seq show an enrichment for FOXJ2 targets by GSEA analysis. **D)** The differentially expressed genes between H3.3K27A cells and control cells as determined by RNA-seq show a depletion for NRSF (REST) targets by GSEA analysis. **E)** Western blot analysis of H3.3K27A and control cells on day 6 of differentiation confirms the increase in REST in the H3.3K27A mutant cells. Lamin B1 is the loading control. **F)** The peak tracks for H3.3K27me3 (pink), H3K27me3 (blue), H3.3 (yellow), and H3 (green) ChIP-seq across the REST gene as well as the RNA-seq expression levels, on day 0 (pluripotency) and day 6 of differentiation, is visualized using UCSC Genome Browser ^98^ (http://genome.ucsc.edu/).

Strikingly, GSEA of the H3.3K27A transcriptome at differentiation also identified a strong depletion of genes regulated by the transcription factor Neuron Restrictive Silencer Factor (NRSF) (Fig. 7D). NRSF, better known as REST (RE1-silencing transcription factor), is important for repressing terminal neuronal genes in non-neuronal cells^67^. REST, a GSEA predicted FOXJ2 target enriched in the H3.3K27A mutants at day 6 of differentiation, exhibits increased expression in the H3.3K27A cells compared to the control, which was validated via WB (Fig. 7A, E, Fig. S12D). ChIP-seq and RNA-seq analysis of mESC shows that REST exhibits high levels of H3.3 at its promoter region in pluripotency, which is correlated with expression of REST, allowing it to transcriptionally repress its target neuronal genes (Fig. 7F). At day 6 of differentiation, there is an increase in H3.3K27 methylation at the promoter of REST, as well as a decrease in overall H3.3 levels. These trends correlate with decreased expression of REST, which may then allow for the expression of terminal neuronal genes during this stage of differentiation (Fig. 7F). Thus, the upregulation of REST in the H3.3K27A cells and subsequent depletion of REST neuronal target genes at day 6 of differentiation may be due to the loss of H3.3K27me3 at REST, resulting in proteomic dysregulation of terminal neuronal genes.

## Discussion

In this work we applied a combination of mass spectrometry and sequencing to profile H3.3K27me3 during stem cell pluripotency and differentiation, gaining critical insights as to the influence of this histone variant modification. SILAC-coupled MS has been used to quantify histone PTM dynamics and inform the chromatin accessibility of a particular PTM^28, 55^. These studies of methylation dynamics determined that histone methylation marks associated with active genes have faster isotope incorporation rates than those associated with repressed genes^28^. Thus, we aimed to determine if H3.3K27 methylation behaves as a repressive or active mark, and found that the incorporation of isotope-labeled methyl groups into H3.3K27me3 is similar to that of known repressive marks. This data suggests that H3.3K27me3 may lie in heterochromatic regions. With high resolution MS we demonstrated that H3.3K27me1 gains two additional methyl groups during differentiation to become H3.3K27me3, which appears in the MS data with two labeled methyl groups (H3.3K27me3^**^). H3.3K27me3^**^ does not reach the same level as H3K27me3^**^ over differentiation, suggesting H3.3K27me3 may occur in more static chromatin environments that are less affected by differentiation, different from H3K27me3. H3.3 has been detected at heterochromatic regions, such as telomeres, as well as at distal intergenic regions^10, 13^. Thus, these relatively compact chromatin regions may host H3.3K27me3. Methylation dynamics have been argued to be influenced solely by histone turnover^54^. However, while this correlation remained true in pluripotent mESCs and active marks in our data, this is not apparent for repressive marks in differentiated cells, suggesting that turnover of these modified histones is not the primary determinant for the regulation of gene expression by these PTMs. Moreover, the quantification of steady-state levels and dynamics indicate that H3.3K27 methylation is maintained over differentiation comparatively to H3K27 methylation.

To explore the biological implications of this maintenance, we performed ChIP-seq for H3.3K27me3 utilizing a novel antibody that preferentially recognizes H3.3K27me3. Strikingly, we find that H3.3K27me3 does not show the same enrichment at the TSS of lowly expressed genes as is observed for H3K27me3, though there was a minor enrichment at the TSS of repressed genes in pluripotency. Furthermore, H3.3K27me3 does not show the same negative correlation as H3K27me3, as would be expected for a repressive mark. Yet, H3.3K27me3 also does not display the same strong positive correlation with gene expression as is exhibited by H3.3 overall^8^. This potentially suggests that H3.3K27me3 acts to neutralize the activating potential of H3.3. Thus, a gene harboring H3.3K27me3, when demethylated, is now marked by H3.3 and is conceivably accessible for transcription. H3.3 has been shown to impair higher order chromatin structures, which is particularly important at the enhancer and promoter regions of RAR/RXR targeted genes that are responsive to retinoic acid^68^. This indicates that both the presence of the histone variant and the modification play a role in gene regulation, unique from that of H3K27me3.

Interestingly, H3.3K27me3 exhibits distinct genomic localization from H3K27me3. While canonical H3K27me3 is enriched at promoter regions in both pluripotency and day 6 of differentiation, H3.3K27me3 is mainly found at promoter regions in pluripotent cells compared to distal intergenic regions in differentiated cells. This suggests that the H3.3K27me3 sites that are maintained over differentiation, as observed in our quantitative analysis, possibly occur in distal intergenic regions. H3.3 has been identified at intergenic regions in both mESC and neuronal precursor cells^13^, and our data now suggests this H3.3 may be marked with K27me3. Intergenic regions possess mainly inaccessible chromatin^69, 70^, consistent with our studies of methylation dynamics showing that H3.3K27me3 may occur in static chromatin regions. Although we find H3.3K27me3 at distal intergenic regions, analysis of our H3.3K27me3 data and publicly available data^58, 71, 72^, suggests these are not enhancers. Thus, the function of H3.3K27me3 at distal intergenic regions has yet to be determined and requires more exploration.

We also determined that in differentiation, H3.3K27me3 displays a unique biological processes enrichment compared to H3K27me3. At this stage, H3K27me3 remains associated with genes involved in development and differentiation, but H3.3K27me3 shows enrichment at lineage-specific genes involved in processes such as adhesion, the nervous system, and sensory perception of smell. Thus, the maintenance of H3.3K27me3 compared to H3K27me3 may be due to the importance of H3.3K27me3 for regulation of lineage-specific genes, such as olfactory receptor genes. For the majority of these olfactory receptor genes, H3.3K27me3 is at the promoter region, and these genes are not expressed. In fact, other chromatin accessibility studies have also found that olfactory genes are enriched for inaccessible promoters^73, 74^, and our methylation dynamics data and ChIP-seq on H3.3K27me3 support the same conclusion.

Olfactory receptors (ORs) are G-protein coupled receptors that are expressed in a monogenic and monoallelic fashion on the surface of olfactory neurons in the brain^75, 76^. In order to allow for expression of only one receptor per olfactory neuron, these genes require strict regulation through an interplay of slow chromatin activation and a potential negative feedback signal^75, 77, 78^. OR genes are located in constitutive heterochromatic regions, marked with H3K9me3 and H4K20me3. One mechanism of regulation proposed silencing through these marks, de-repression of which would then allow for a low expression level of a chosen OR gene^79^. While there has been no evidence so far that OR genes are marked with H3K27me3, H3.3 has been described at constitutive heterochromatin through deposition by the chaperones ATRX/DAXX, coinciding with the H3K9me3 mark^13, 80^. Our data is the first to describe H3.3K27me3 at these olfactory receptor genes. Interestingly, we do not detect an increase in OR gene expression in the H3.3K27A mutant cells. However, this is not completely surprising given that these genes are also marked with H3K9me3 and H4K20me3, which are unaffected in the H3.3K27A mutant cells. OR genes require precise regulation, and thus loss of one histone PTM may not be enough to cause dysregulation. The details of this regulation of OR genes by H3.3K27me3 remain to be defined by future work.

Since we identified unique characteristics of H3.3K27me3 compared to H3K27me3, we sought to interrogate the effect of the loss H3.3K27 methylation in the context of this early developmental transition. A limitation of the H3.3K27A mutant cells is that these mutants lack not only methylation on H3.3K27, but acetylation as well. Thus, although it may not be possible to associate phenotypic changes with loss of either mark, these cells are still a critical tool to understand the importance of H3.3K27 in differentiation. In agreement with previous studies describing only minor changes in the transcriptome of H3.3K27R mESC in the pluripotent state^26^, we also observe few changes in the proteome and overall histone PTM landscape in the H3.3K27A mESC during pluripotency. Alterations were more evident at later stages of differentiation, suggesting a critical role for this residue during cellular differentiation.

The interplay between histone PTMs has been well-documented^59^. Notably, we find that some histone PTM changes that are observed in the H3.3K27A mutants at day 6 of differentiation are reflective of the histone PTM changes observed in H3.3K27M gliomas^81^. Specifically, the H3.3K27A mutants exhibit a global increase in H3K27ac and a global decrease in H3K27me3. H3K27ac is an important enhancer mark, while H3K27me3 is involved in repression. Additionally, the H3.3K27A mutants exhibit an increase in another well-characterized mark, H3K4me3, which is enriched at promoter regions and associated with transcriptional activation. Thus, H3.3K27 demonstrates a critical role in maintaining the balance with other regulatory histone PTMs during differentiation, and the disruption in the histone PTM landscape results in the large proteomic changes that are observed. Interestingly, not only did we detect PTM changes associated with H3.3K27M gliomas in our H3.3K27A differentiating cells, but we also identified that H3.3K27A cells at this stage exhibit a significant increase in proteins involved in glial differentiation. This indicates that H3.3K27 is necessary for proper glial differentiation through the influence of several histone PTMs. Although the H3.3K27A mutant results in a similar decrease in global levels of H3K27me3, the mechanism of this decrease most likely differs from what occurs in H3.3K27M gliomas. In gliomas, the H3.3K27M mutation exerts a dominant negative effect, in which the H3.3K27M peptide allosterically inhibits EZH2, the methyltransferase responsible for H3K27me3^82^. Given that the most potent inhibition of EZH2 occurred with amino acids with long, hydrophobic side chains and minimal branching^82^, as is the case for methionine and isoleucine, it is unlikely that the K27A mutation acts through the same mechanism.

In both pluripotency and differentiation, we detect a large increase in H3.3K36me3 in the H3.3K27A mutant cells compared to the control cells. This is expected, given previous studies demonstrating an antagonistic effect between H3K27me3 and H3K36me3^60, 61^. In fact, H3K36me3 may directly inhibit the writing of H3K27me3 by PRC2^62^. H3K36me3 correlates with transcriptional elongation and is found across the bodies of actively transcribed genes^83^. Studies in yeast have found that H3K36me3 recruits histone deacetylases (HDACs) to remove acetylation marks on H3 and H4 and prevent aberrant initiation of transcription^84, 85^. Methylation on H3.3K36 has also been suggested to recruit MOF, the acetyltransferase for H4K16ac, in neuronal stem cells to allow for proper neuronal differentiation^14^. Consistent with these studies, we detect a global decrease in H4 acetylation in the H3.3K27A mutant cells during differentiation, with only H4K16ac being increased in the H3.3K27A cells. This demonstrates the direct interplay between H3.3K27 and other histone PTMs that ensure proper epigenetic regulation and cell differentiation.

Interestingly, our work has identified potential correlations between H3.3K27me3 and TFs important for embryonic stem cell differentiation. We identified an enrichment of H3.3K27me3 at FOXJ2/3 motifs during differentiation. Notably, in differentiated H3.3K27A mutant cells, we found increased expression of FOXJ2/3 targets at the RNA and protein levels through GSEA. Both FOXJ2 and FOXJ3 have been implicated in neural development, exhibiting expression in the neuroectoderm and subsequently the notochord, neural tube, and neural crest in the developing embryo^63, 86^. Although both have been suggested to be transcriptional activators^65, 66^, their exact biological roles are not yet fully understood. FOXJ2 over-expression has been suggested to have a negative effect on development based on embryonic lethality in mice^87^. Likewise, FOXJ3 has been characterized to prevent mESC from differentiating into neural cells, either through repression or activation of downstream targets^88^. In our work, we identify that H3.3K27me3 is enriched for FOXJ2/3 binding motifs during stem cell differentiation, and upon loss of methylation at H3.3K27, as is the case in the H3.3K27A cells, there is an upregulation of FOXJ2/3 targets, suggesting a potential regulatory role for H3.3K27me3 at these motifs during stem cell differentiation. As transcriptional activators, it is consistent that we observe increased expression of FOXJ2/3 targets in H3.3K27A cells, where their binding motif is presumably more accessible due to loss of H3.3K27me3. Curiously, FOXJ2 has been associated with the regulation of olfactory receptor genes since its binding motif was found at enhancers of these genes^77^, suggesting another potential link between H3.3K27me3 and the regulation of lineage specific genes.

One of the putative FOXJ2 targets is REST, a transcriptional repressor of terminal neuronal genes. REST protein expression was significantly increased in the H3.3K27A cells compared to the control cells during differentiation. Consistent with this increase in REST, we detected a strong depletion of REST targets at the transcript level in H3.3K27A cells during differentiation. In pluripotent stem cells and neuronal precursor cells, REST represses neuron-specific genes involved in synaptogenesis, synaptic plasticity and remodeling, synaptic vesicle maturation, and adhesion proteins^89-92^. Similarly, H3.3K27me3 during differentiation was enriched at genes involved in cell adhesion and the positive regulation of synapse assembly, supporting the connection between H3.3K27me3 and TFs involved in proper neuronal differentiation. One of the mechanisms by which REST acts is through epigenetic remodeling via recruitment of HDACs^93^. We detected a global decrease in H4 acetylation in the H3.3K27A cells during differentiation, which is expected since an increased level of REST in these cells may lead to increased recruitment of HDACs to target genes. Whether PRC2 directly interacts with REST has been debated, with some studies suggesting a co-regulatory role for these complexes in neuronal differentiation^94, 95^, while other studies indicate the existence of distinct regulatory pathways^96^. However, previous studies have not differentiated between H3K27me3 and H3.3K27me3, nor explored direct regulation of REST expression by histone methylation. REST is expressed in mESC during pluripotency, but dramatically decreases in expression during differentiation, which is important for the acquisition of the neuronal phenotype^97^. Expression of REST in pluripotency is correlated with low levels of H3.3K27me3, which increase upon differentiation. The H3.3K27A mutant cells have complete loss of methylation on H3.3K27, and exhibit a failure to repress REST expression during differentiation, leading to increased expression of REST and decreased expression of REST targets. Thus, H3.3K27me3 may play a role in ensuring proper timing of differentiation through the regulation of TFs of terminal neuronal genes as well as lineage-specific genes themselves.

## Conclusions

Using a multi-omics profiling approach, we find that methylation on the variant H3.3K27 differs from canonical H3K27me3 with respect to its overall levels and genomic localization during cellular differentiation. We report that H3.3K27me3 is uniquely enriched at promoter regions of lineage-specific genes, such as olfactory receptors, and is potentially involved in the regulation of the TFs FOXJ2/3 and REST, which are necessary for proper differentiation to the neuronal lineage. This suggests a distinct but critical role for H3.3K27me3 during embryonic stem cell differentiation. Future work should aim to explore the regulatory nature of H3.3K27me3 at FOXJ2/3 binding sites, including REST. Additionally, details on the role of H3.3K27me3 in olfactory receptor gene regulation remains to be elucidated. Whether H3.3K27me3 exhibits similar enrichment when differentiating to other cell lineages remains to be seen. Interestingly, beyond enrichment at promoter regions, H3.3K27me3 is more highly enriched at distal intergenic regions in differentiating cells. However, questions remain as to the biological significance of H3.3K27me3 at these regions. Overall, our work reports several potential roles of H3.3K27me3 during stem cell differentiation, and highlights the importance of differentiating between modifications on canonical histones versus their variants, as they may exhibit different characteristics and biological functions.

## Author Contributions

B.A.G. conceived and designed the project. Y.K. performed the experiments. M. T. generated the CRISPR cell lines under the supervision of K.M.N. Then Y.K., P.J.L., S.S. and Z.F.Y. analyzed the data. Y.K. and P. J. L. interpreted the data and generated the figures. Y.K. wrote the paper. B.A.G. provided supervision. All authors reviewed the manuscript.

## Acknowledgments

We thank members of the Garcia lab for scientific input. This work was supported by grants from the National Institutes of Health: 5T32GM071339-13, 2T32CA009140-41A1, CA196539, NS111997, and AG031862. Additionally, we acknowledge support from: St. Jude Children’s Research Hospital Collaborative on Chromatin Regulation in Pediatric Cancer, and the Crohn’s and Colitis Foundation (RFA598467). We also gratefully acknowledge funding from the Blavatnik Family Foundation.

