## Supplemental Methods and Figures for "Multi-omic profiling of histone variant H3.3 lysine 27 methylation reveals a distinct role from canonical H3 in stem cell differentiation"

Supplementary Methods
Supplementary Figures: 13

#### Supplementary Methods

*Histone sample preparation*

Nuclei were extracted from the collected cell pellets using nuclei isolation buffer (NIB) as previously described[1]. Briefly, cell pellets were first washed with NIB, and then lysed using 10 cell pellet volumes of NIB + 0.2% NP-40. Subsequently two washes with NIB without NP-40 were performed, to remove any trace of detergent. Once nuclei were isolated, histones were extracted with 0.2 M $H_2SO_4$, followed by precipitation with a final concentration of 33% trichloroacetic acid (TCA) overnight on ice in the cold room. Precipitated histones were washed with ice-cold acetone to remove the acid. The derivatization and digestion of the histones was also performed as previously described[1], with minor adjustments. Histones were derivatized with propionic anhydride for two rounds prior to trypsin digestion overnight (enzyme:sample 1:20), followed by two more round of propionylation.

*Metabolite analysis*

Metabolites were extracted as previously described[2], by resuspending cell pellets in 1mL of cold (-80°C) solution of 80% methanol/20% water, and then incubating in -80°C freezer for 15 minutes, followed by centrifugation and collection of the supernatant. This process was repeated for a second extraction and both supernatants were combined, dried, and stored at -80°C until MS analysis.

Metabolite samples were analyzed by LC-MS with a Vanquish UHPLC (Thermo Fisher Scientific) coupled to a QQQ mass spectrometer (Thermo Fisher Scientific) operating in positive mode. A Hypersil Gold C18 reverse phase column was used. The 10 minute HPLC gradient was: hold at 1% B for 2 min (A = 0.1% formic acid; B = 80% acetonitrile, 0.1% formic acid), from 1% B to 40% B within 4 min, up to 95% B in 0.1 min and hold at 95% B for 1.9 min, then back down to 1% B in 0.1 min and hold at 1% B for 1.9 min. The flow rate was 300 ul/min. The column was heated to 45°C. Data were acquired using a selected reaction monitoring (SRM) experiment. For the unlabeled SAM, the precursor m/z targeted was 399.052, and fragment ions 135.946, 250, 264.012, and 298.054 m/z. For the $^{13}CD_3$-SAM the precursor m/z targeted was 403.14, and fragment ion 250 m/z. The dwell time was 10 ms. The Q1 resolution (FWHM) was 0.7, Q3 resolution (FWHM) was 1.2, and CID gas at 1.5 mTorr. For MS analysis, the dried metabolites were resuspended in 100ul MS-grade water, and 1 ul was used for injection. An unlabeled purified SAM standard was also used at 1 pmol per injection, as a control for retention time alignment.

The SRM QQQ data was analyzed manually using Xcalibur to extract for the precursor and fragment masses. The quantitation was performed by taking the AUC for the 250 m/z fragment corresponding to the unlabeled SAM and to the $^{13}CD_3$-SAM. The labeling ratio was calculated by taking the aforementioned AUC of the $^{13}CD_3$-SAM as the numerator, and the denominator was the sum of the AUC from the unlabeled SAM plus the $^{13}CD_3$-SAM. The averages and standard deviations were calculated in Microsoft Excel. To normalize the $^{13}CD_3$-methionine incorporation histone data with the SAM labeling data, the fold change between the SAM labeling ratio in pluripotent mESC vs differentiating cells was calculated at each metabolite time point. The ratio of heavy methylation incorporation was re-calculated for the differentiating cells using the normalized AUCs.

*Salt fractionation*

Salt fractionation was performed as previously described[3]. Briefly, $20*10^6$ cells were lysed in buffer 1 (0.32 M sucrose, 60 mM KCl, 15 mM NaCl, 5 mM $MgCl_2$, 0.1 mM EGTA, 15 mM Tris pH 7.5, 0.5 mM DTT, 0.1 mM PMSF) and then the nuclei were layered onto buffer 2 (1.2 M sucrose, 60 mM KCl, 15 mM NaCl, 5 mM $MgCl_2$, 0.1 mM EGTA, 15 mM Tris pH 7.5, 0.5 mM DTT, 0.1 mM PMSF). The nuclei were pelleted and resuspended in buffer 3 (10 mM Tris pH 7.4, 2 mM $MgCl_2$, HALT protease inhibitors) and supplemented with 5 mM $CaCl_2$, and the DNA was digested to mono-nucleosomes by adding MNase. After centrifugation, the resulting pellet was fractionated in varying concentrations of salt (NaCl = 75 mM, 150 mM, 300 mM, 600 mM) in buffer 4 (10 mM Tris pH 7.4, 2 mM $MgCl_2$, 2 mM EGTA, 0.1% Triton X-100).

*Western blotting and dot blotting*

Antibodies for ChIP-seq were assessed by dot blot and WB. For dot blot, 1 ug of purified peptide (H3.3K27me3, H3K27me3, unmodified H3.3K27, H3K9me3) was spotted onto a 0.2 um nitrocellulose membrane, and blocked for 1 hour at room temperature in 5% milk in Tris buffered saline (TBS) supplemented with 0.1% Tween 20 (TBST). Primary antibodies were diluted 1:1000 in 5% milk in TBST and incubated overnight at 4°C. The membrane was washed three times with TBST followed by incubation with HRP-conjugated secondary antibody for 1 hour at room temperature in 5% milk/TBST. The membrane was washed again and imaged by Fujifilm LAS-4000 imager.

For WBting, after ChIP, the beads were resuspended in 4X LDS Sample buffer (NuPAGE, Invitrogen) + 2% β-mercaptoethanol (Biorad). Samples were boiled at 95°C and run on a 12% Bis-Tris gel (NuPAGE). After transfer to nitrocellulose membrane, the blocking, blotting, and washing was performed as described above.

For the αREST WB, nuclear extraction of the day 0 and day 6 samples was prepared as previously described[4]. Briefly, nuclear extracts were prepared by lysing mESC in buffer A (10mM HEPES pH 7.9, 1.5mM $MgCl_2$, 10mM KCl) supplemented with protease inhibitors on ice, followed by centrifugation. Cell pellets were further homogenized in buffer A supplemented with 0.15% NP-40 and protease inhibitors. The nuclei were pelleted, then nuclear proteins were released for 1 hour at 4°C in Buffer C (420mM NaCl, 20mM HEPES pH 7.9, 20% glycerol, 2mM $MgCl_2$, 0.1% NP-40 and protease inhibitors), and the supernatant was saved as the soluble lysate. The insoluble chromatin pellet was further digested in nuclear pellet solubilization buffer (150 mM NaCl, 50 mM Tris pH 8.0, 1% NP-40, and 5 mM $MgCl_2$) supplemented with Benzonase and incubated at room temperature, shaking, for 10 minutes. After centrifugation, the supernatant was combined with the previous soluble lysate. Protein concentration was estimated by Bradford assay. WB was conducted using a 4-12% Bis Tris gel (NuPAGE).

*qPCR for gene expression analysis*

Cells were homogenized using QIAshredder (Qiagen), and RNA was extracted using a RNAeasy kit (Qiagen 74014) following the manufacturer's guidelines. cDNA was synthesized using a Superscript III

First Strand Synthesis System (Thermo Scientific 18080051). qPCR was performed using the Power SYBR Green PCR Master Mix (Applied Biosystems) and the Real-Time PCR System (Applied Biosystems) following the manufacturer's instructions. The qPCR primers used for Nanog, Oct4, Gata4, Cyclin D1, and GAPDH were the same as previously published[5]. The relative abundance of mRNA was calculated as a quantity mean, and the markers Nanog, Oct4, Gata4, and Cyclin D1 were each normalized to the quantity mean of the housekeeping gene GAPDH.

*Proteomics sample preparation*

Proteomics samples were prepared following ProtiFi s-trap protocol[6]. Cells were lysed in 1x SDS lysis buffer (5% SDS in 100 mM TEAB, pH 7.55), and DNA was sheared by probe sonication. 50 ug of proteins were reduced with dithiothreitol (DTT) at a final concentration of 20mM at 95°C for 10 minutes, and then alkylated with iodoacetamide to a final concentration of 40 mM and incubated 30 minutes in the dark. Aqueous phosphoric acid was then added to a final concentration of 1.2%, and then six volumes of s-trap buffer (90% methanol,100 mM TEAB, pH 7.55). S-traps were made in-house using a P200 pipette tip, packed with one 2mm plug of C18 disk, and two 3mm plugs of a quartz disk. Sample was loaded onto the s-trap and washed with s-trap buffer 3 times. Protein digestion was performed on the s-trap column for 1 hour at 47°C in digestion buffer (50mM TEAB) containing trypsin (Promega) at a 1:20 wt:wt (enzyme:protein) ratio. Peptides were eluted using three buffers: first 50 mM TEAB, then 0.2% formic acid, and lastly 50% acetonitrile containing 0.2% formic acid. Samples were dried in a speed-vac and desalted by stage-tipping[1].

### Supplementary Figures

A

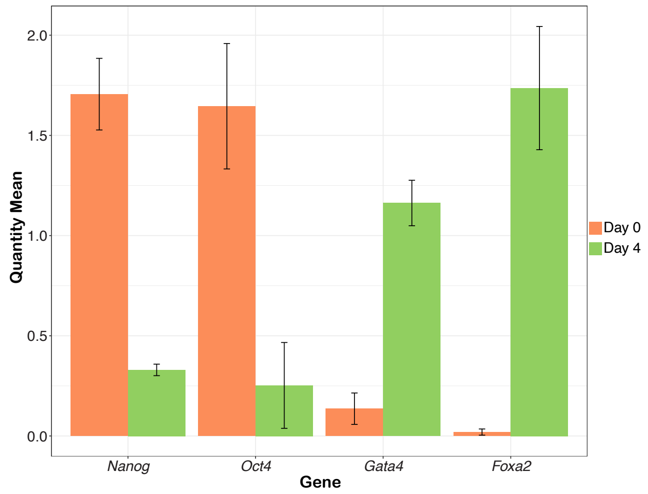


B

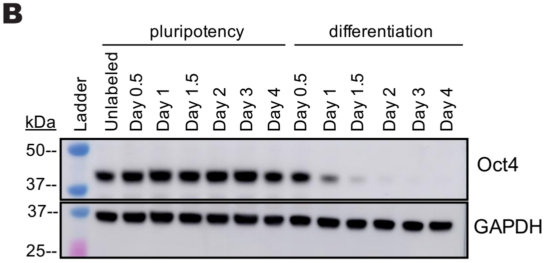


C

| Synthetic Peptide | Sequence |
|---|---|
| H3.3 27-40 unmodified | KSAPSTGGVKKPHR |
| H3.3K27me1 | **Kme1**SAPSTGGVKKPHR |
| H3.3K27me2 | **Kme2**SAPSTGGVKKPHR |
| H3.3K27me3 | **Kme3**SAPSTGGVKKPHR |
| H3 27-40 unmodified | KSAPATGGVKKPHR |
| H3K27me1 | **Kme1**SAPATGGVKKPHR |
| H3K27me2 | **Kme2**SAPATGGVKKPHR |
| H3K27me3 | **Kme3**SAPATGGVKKPHR |

D

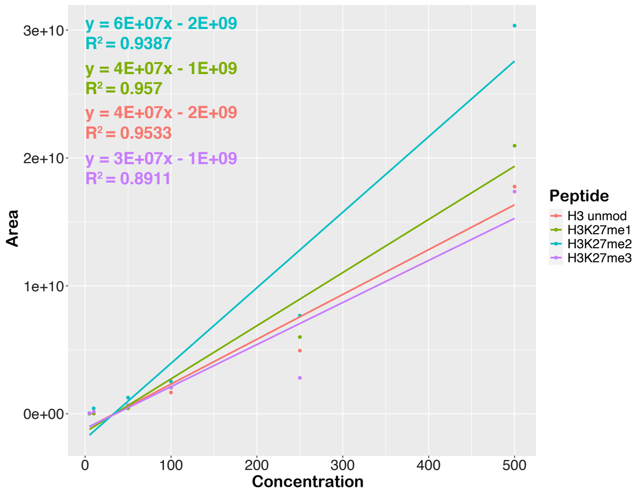


E

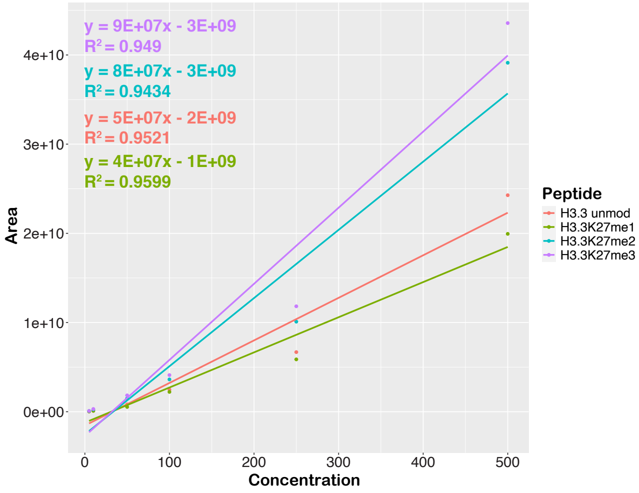

**Supplemental Figure 1. Using isotopically labeled synthetic histone peptides to generate standard curves allows for absolute quantification of endogenous histone peptides. A)** RT-qPCR of mouse embryonic stem cells (mESC) in pluripotency (day 0) and day 4 of differentiation, analyzing for pluripotency markers *Nanog* and *Oct4*, and differentiation markers *Gata4* and *Foxa2*. Samples were normalized to the *Gapdh* control, and the quantity mean is plotted, with the quantity standard deviation indicated (technical triplicates). **B)** Western blot analysis of whole cell lysate from mESC at different time points in pluripotency and differentiation, blotting for the pluripotency marker Oct4 (39 kDa) with Gapdh (35.8 kDa) as the loading control. **C)** The isotopically labeled synthetic peptide sequences (Cell Signaling Technology) are listed. The bolded residue indicates the lysine that is post-translationally modified with either mono-methylation (me1), di-methylation (me2), or tri-methylation (me3). The heavy isotope labeled residues are indicated in red. The isotopically labeled proline (P) increases the normal isotope peptide mass by 6.013805. The isotopically labeled valine (V) plus the isotopically labeled proline (P) increase the normal isotope peptide mass by 12.02761. The isotopically labeled alanine (A) plus the isotopically labeled proline (P) increase the normal isotope peptide mass by 10.0209. **D)** Standard curve of the H3 synthetic peptide isotopically labeled standards (listed in panel C). The concentrations of these standards analyzed by LC-MS were: 500 femtomole (fmol), 250 fmol, 100 fmol, 50 fmol, 10 fmol, and 5 fmol. Standards were analyzed in technical duplicates, with the area under the curve (AUC) for each concentration and peptide plotted. The best fit line for each peptide, as determined in Microsoft Excel, is plotted, along with the equation for the best fit line. **E)** Standard curve of the H3.3 synthetic peptide isotopically labeled standards (listed in panel C). The standard curve is plotted the same as described in D).

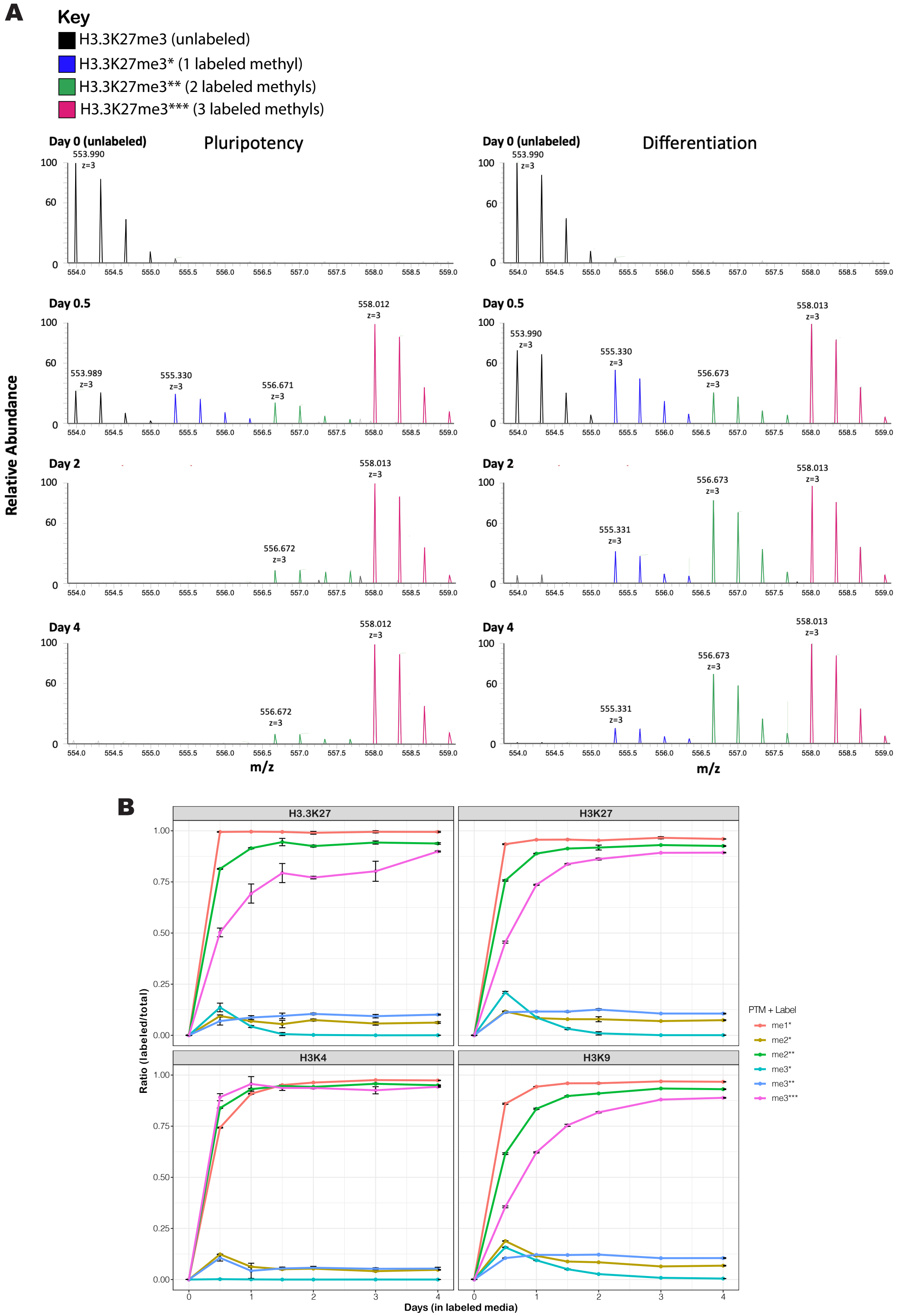
A
Key
H3.3K27me3 (unlabeled)
H3.3K27me3* (1 labeled methyl)
H3.3K27me3** (2 labeled methyls)
H3.3K27me3*** (3 labeled methyls)
Pluripotency
Differentiation
Day 0 (unlabeled)
Day 0.5
Day 2
Day 4
Relative Abundance
m/z
B
H3.3K27
H3K27
H3K4
H3K9
Ratio (labeled/total)
Days (in labeled media)
PTM + Label
me1*
me2*
me2**
me3*
me3**
me3***

**Supplemental Figure 2. $^{13}CD_3$-methionine labeling of histone methylation is detected across time points in pluripotency and differentiation. A)** The isotopic pattern for H3.3K27me3 peptide (+3 charge state) shows incorporation of the heavy labeled methyl groups (resulting in a +1.33 Da mass shift per heavy methyl group, since it is the +3 charge state of the peptide) across the time points in both pluripotency and differentiation. **B)** Heavy methylation incorporation ($^{13}CD_3$-methionine) is plotted over time points in labeling media for the histone marks H3.3K27 (top left), H3K27 (top right), H3K4 (bottom left) and H3K9 (bottom right) during mESC pluripotency. Time points collected were at 0 (unlabeled control), 0.5, 1, 1.5, 2, 3, and 4 days after switching to labeled media. The ratio of heavy labeling was calculated by taking the AUC for a specific heavy labeled methylated species as the numerator, and the denominator is the sum of the AUCs for all labeled and unlabeled forms of that methylated species. The number of asterisks indicate the number of methyl groups that are heavy labeled. The heavy labeling ratios at each time point are averages of biological triplicates, and the standard deviation of the mean is indicated.

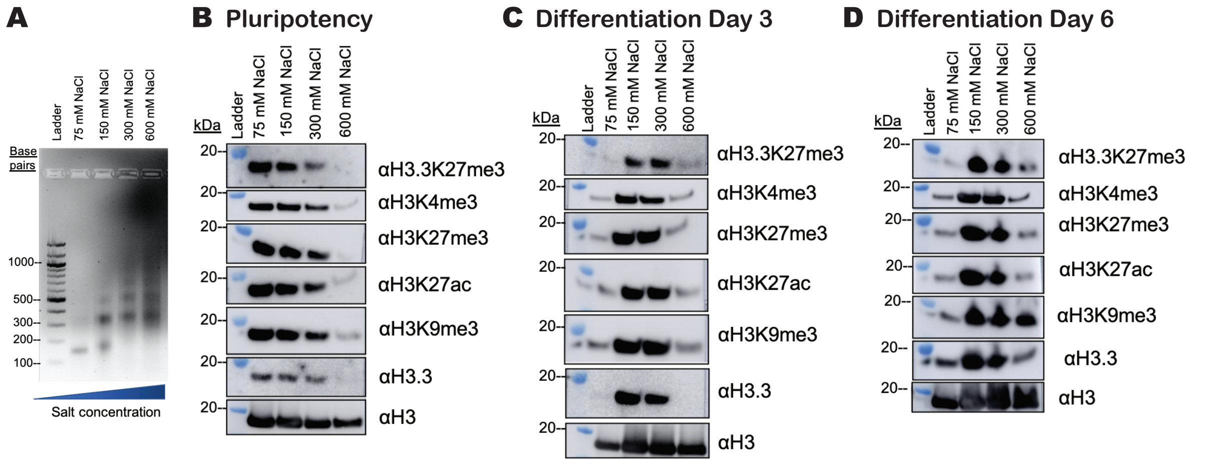


**Supplemental Figure 3. Histone PTM interrogation of chromatin from salt fractionation indicates changes in chromatin compaction across pluripotency and differentiation. A)** 500 nanograms of DNA extracted from salt fractionation samples was run on a 2% agarose gel, indicating mono- (~147 bp), di- (~294 bp), and tri- (~441 bp) nucleosomes extracted with increasing salt concentration. **B)** Western blot analysis of salt fractionation samples of pluripotent mESC, blotted for several histone post-translational modifications, shows that in pluripotency chromatin is more open overall and is extracted at the lower salt fractions. Histone H3 is the loading control. **C)** Western blot analysis performed as described in panel B, but for differentiating mESC at the day 3 time point, shows that chromatin becomes more compact, and extraction shifts overall to higher salt fractions. **D)** Western blot analysis performed as described in panel B, but for differentiating mESC at the day 6 time point, shows that chromatin is extracted at the higher salt fractions, especially H3K9me3 which is found at 600 mM NaCl, as expected due to its enrichment in heterochromatin.

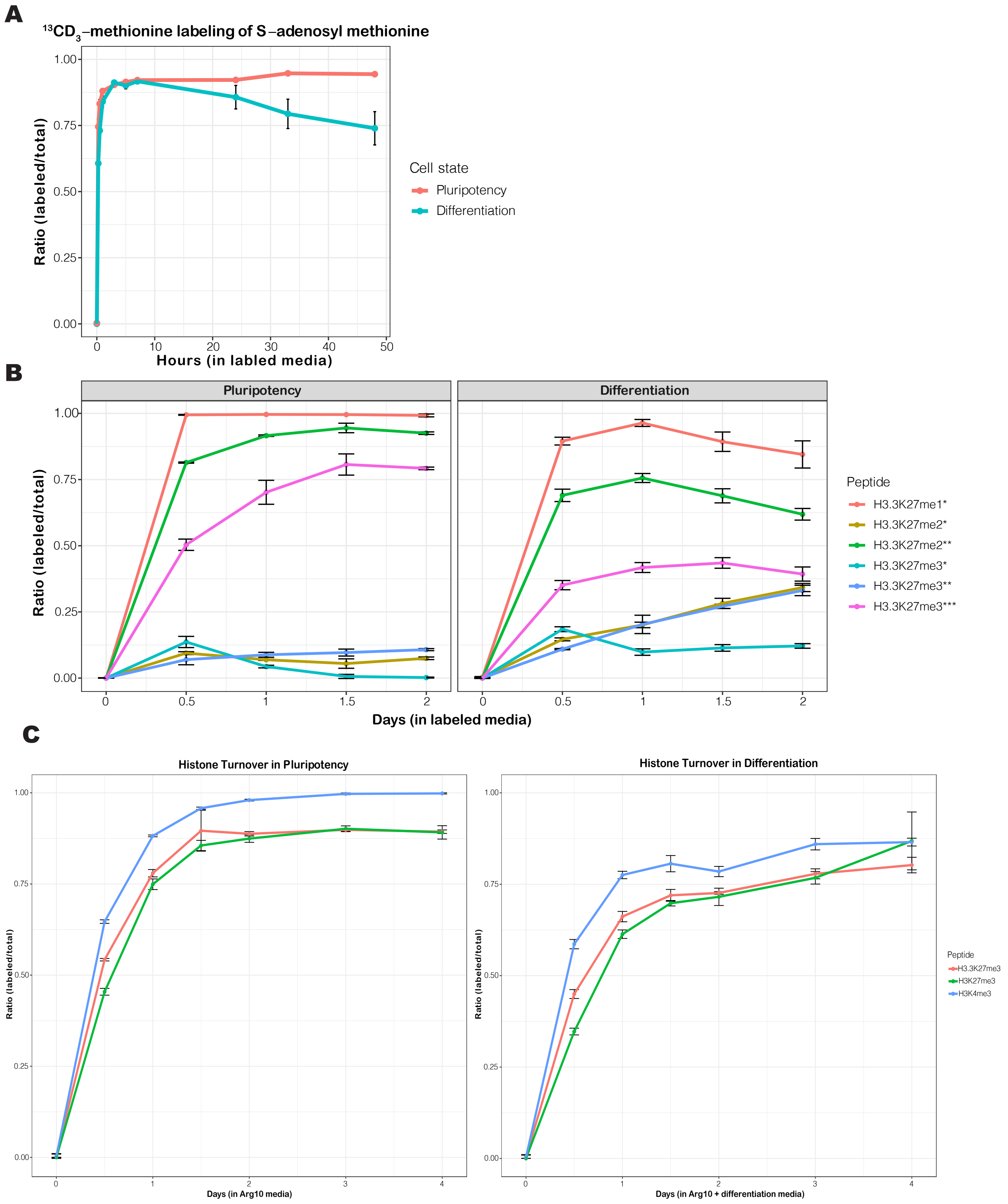
A
13CD3–methionine labeling of S–adenosyl methionine
Ratio (labeled/total)
Hours (in labled media)
Cell state
Pluripotency
Differentiation
B
Pluripotency
Differentiation
Ratio (labeled/total)
Days (in labeled media)
Peptide
H3.3K27me1*
H3.3K27me2*
H3.3K27me2**
H3.3K27me3*
H3.3K27me3**
H3.3K27me3***
C
Histone Turnover in Pluripotency
Histone Turnover in Differentiation
Ratio (labeled/total)
Days (in Arg10 media)
Days (in Arg10 + differentiation media)
Peptide
H3.3K27me3
H3K27me3
H3K4me3

**Supplemental Figure 4. H3.3K27me3 dynamics between pluripotency and differentiation are independent of metabolite cofactor labeling and histone turnover.** A) Heavy methylation labeling of SAM is plotted over the $^{13}CD_3$-methionine labeling time points for both pluripotent (red) and differentiating (blue) cells. Time points collected: 0 hours (unlabeled control), as well as 15 min, 30 min, 1 hour, 3 hours, 5 hours, 7 hours, 24 hours, 33 hours, 48 hours after switching to labeled media. The ratio was calculated by taking the AUC of the labeled SAM over the sum total of the AUC of unlabeled plus labeled SAM. **B)** Heavy methylation labeling ($^{13}CD_3$-methionine) was normalized for the differences in SAM labeling as determined in panel A, and plotted over labeling time points in both pluripotency and differentiation, to allow for comparison of histone methylation incorporation between the two cell states. **C)** The ratio of $^{13}C_6$,$^{15}N_4$-arginine labeling (Arg10) for three histone peptides: H3K27me3 (green), H3K4me3 (blue), and H3.3K27me3 (red), was plotted across Arg10 labeling time points in both pluripotency and differentiation to interrogate histone turnover in the two cell states.

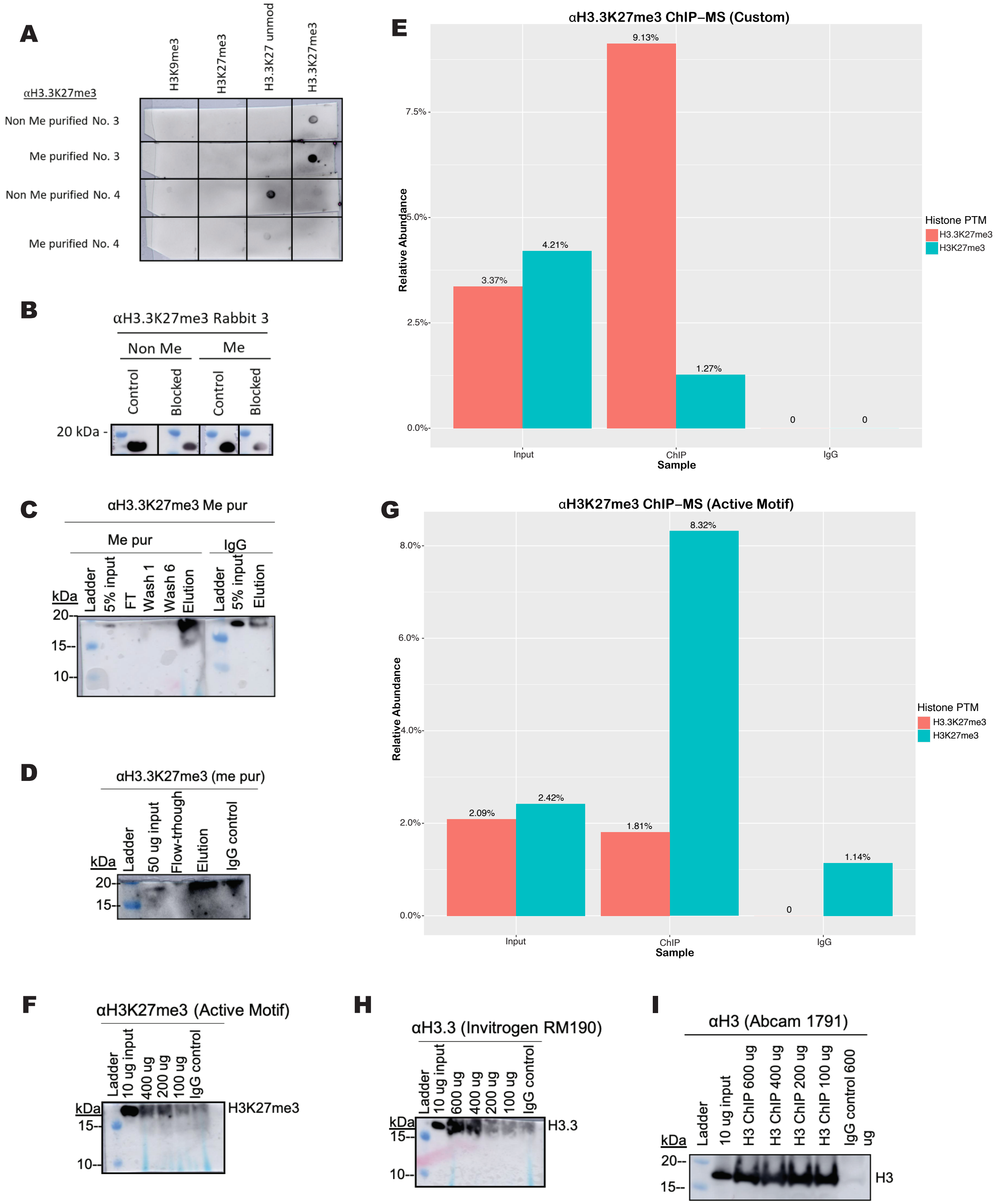
A
αH3.3K27me3
H3K9me3
H3K27me3
H3.3K27 unmod
H3.3K27me3
Non Me purified No. 3
Me purified No. 3
Non Me purified No. 4
Me purified No. 4
B
αH3.3K27me3 Rabbit 3
Non Me
Me
Control
Blocked
Control
Blocked
20 kDa -
C
αH3.3K27me3 Me pur
Me pur
IgG
kDa
Ladder
5% input
FT
Wash 1
Wash 6
Elution
Ladder
5% input
Elution
20-
15-
10-
D
αH3.3K27me3 (me pur)
kDa
Ladder
50 ug input
Flow-trhough
Elution
IgG control
20-
15-
E
αH3.3K27me3 ChIP–MS (Custom)
Relative Abundance
0.0%
2.5%
5.0%
7.5%
3.37%
4.21%
9.13%
1.27%
0
0
Input
ChIP
IgG
Sample
Histone PTM
H3.3K27me3
H3K27me3
G
αH3K27me3 ChIP–MS (Active Motif)
Relative Abundance
0.0%
2.0%
4.0%
6.0%
8.0%
2.09%
2.42%
1.81%
8.32%
0
1.14%
Input
ChIP
IgG
Sample
Histone PTM
H3.3K27me3
H3K27me3
F
αH3K27me3 (Active Motif)
kDa
Ladder
10 ug input
400 ug
200 ug
100 ug
IgG control
15-
10-
H3K27me3
H
αH3.3 (Invitrogen RM190)
kDa
Ladder
10 ug input
600 ug
400 ug
200 ug
100 ug
IgG control
15-
10-
H3.3
I
αH3 (Abcam 1791)
kDa
Ladder
10 ug input
H3 ChIP 600 ug
H3 ChIP 400 ug
H3 ChIP 200 ug
H3 ChIP 100 ug
IgG control 600 ug
20-
15-
H3

**Supplemental Figure 5. Quality control experiments confirm specificity and utility of H3.3K27me3 custom antibody for ChIP-seq. A)** Dot blot analysis of the H3.3K27me3 target peptide, and three potential off-target peptides (H3K9me3, H3K27me3, H3.3K27 unmodified), blotted using purified IgG H3.3K27me3 antibody from two rabbits (No. 3 and No. 4, ProteinTech). "Non Me purified" means the antibody was collected as flow through from the first purification column (bovine serum albumin peptide column), which was used to deplete non-specific antibody binding. "Me purified" means that the antibody elution was collected after the second purification column (H3.3K27me3 peptide column) which was used to enrich for specific antibody binding. **B)** Western blot analysis of a peptide competition assay, where the αH3.3K27me3 antibody was pre-blocked (labeled "blocked") overnight with an 10X excess of target peptide, H3.3K27me3, and then used for western blotting of 5 ug histones. Decreased signal intensity using the blocked antibody compared to the unblocked control antibody indicates specificity for the target peptide H3.3K27me3 that was used for blocking. **C)** Western blot analysis of a mono-nucleosome immunoprecipitation using the custom αH3.3K27me3 antibody and an IgG control antibody. 5% input, flow-through (FT), the first and last wash (wash 1 and wash 6), as well as the elution were analyzed, and indicated successful enrichment in the elution. **D)** Chromatin immunoprecipitation followed by western blot (ChIP-WB) using the custom αH3.3K27me3 antibody and an IgG control antibody. 600 ug of input was used for the ChIP, and for WB analysis 50 ug of input, flow-through, and elution were analyzed and indicated successful enrichment in the elution. **E)** Chromatin immunoprecipitation followed by mass spectrometry analysis (ChIP-MS) using the custom αH3.3K27me3 antibody and the IgG control antibody. ChIP was conducted the same way as described in panel D. The αH3.3K27me3 antibody is observed to preferentially enrich for H3.3K27me3 (red) compared to canonical H3K27me3 (blue). Additionally, the αH3.3K27me3 antibody successfully enriches for H3.3K27me3 compared to the input and IgG control samples. **F)** ChIP-WB of an αH3K27me3 antibody (Active Motif) and IgG control antibody. Various amounts (400 ug, 200 ug, and 100 ug) were tested as input for the ChIP. Although there are limitations with efficiency (low yield), specificity for H3K27me3 is observed. **G)** ChIP-MS of αH3K27me3 antibody (Active Motif) and IgG control antibody using 600 ug of input confirms specificity of the antibody for H3K27me3 (blue) compared to H3.3K27me3 (red). Additionally, the αH3K27me3 antibody successfully enriches for H3K27me3 compared to the input and IgG control samples. **H)** ChIP-WB of αH3.3 antibody (Invitrogen) and IgG control antibody for comparison, where various amounts (600 ug, 400 ug, 200 ug, and 100 ug) were tested as input for ChIP. For ChIP-seq, this H3.3 antibody was used as a control with 600 ug of input. **I)** ChIP-WB of αH3 antibody (Abcam) and IgG control antibody for comparison, where various amounts (600 ug, 400 ug, 200 ug, and 100 ug) were tested as input for ChIP. For ChIP-seq, this H3 antibody was used as a control with 600 ug of input.

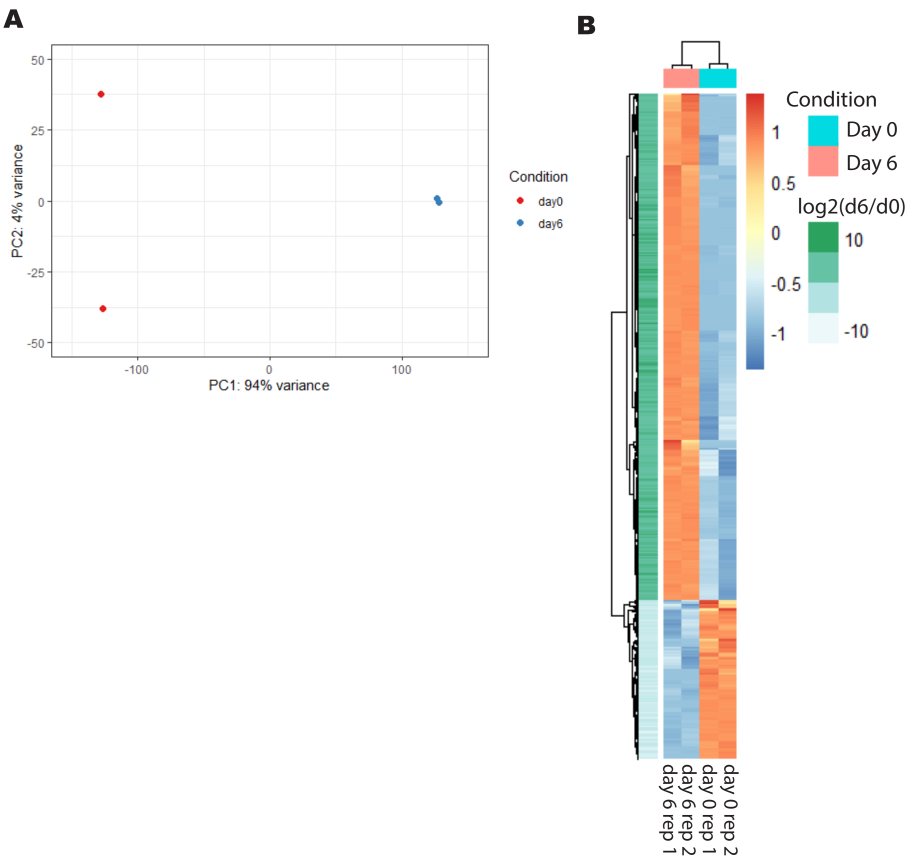


**Supplemental Figure 6. RNA-seq confirms transcriptomic differences between pluripotency (day 0) and differentiation (day 6). A)** Principal component analysis (PCA) of RNA-seq samples shows that biological replicates cluster together, and that the different time points (day 0 = red, day 6 = blue) separate well. **B)** Heatmap of the genes detected in the RNA-seq analysis for both day 0 (turquoise) and day 6 (pink). Biological replicates are plotted separately. The z-score is indicated by the blue – red color scale, and the fold change of day 6 / day 0 for each gene is indicated in the left column using the green color scale. Clear gene expression differences are observed between day 0 and day 6, and biological replicates cluster together.

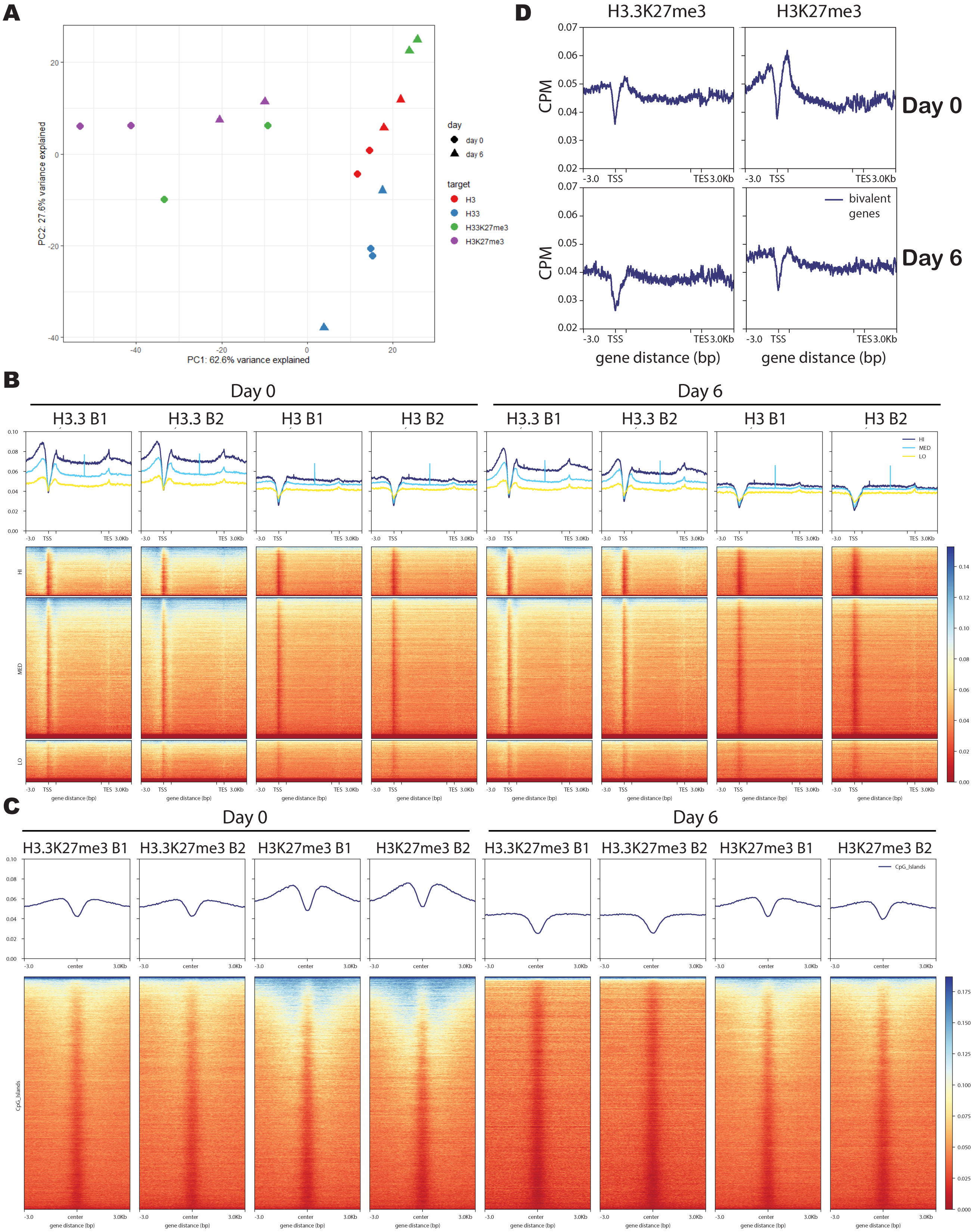
A
PC1: 62.6% variance explained
PC2: 27.6% variance explained
day
day 0
day 6
target
H3
H33
H33K27me3
H3K27me3
D
H3.3K27me3
H3K27me3
CPM
Day 0
Day 6
bivalent genes
gene distance (bp)
B
Day 0
Day 6
H3.3 B1
H3.3 B2
H3 B1
H3 B2
HI
MED
LO
gene distance (bp)
C
Day 0
Day 6
H3.3K27me3 B1
H3.3K27me3 B2
H3K27me3 B1
H3K27me3 B2
CpG_Islands
gene distance (bp)

**Supplemental Figure 7. H3.3K27me3 ChIP-seq correlation with gene bivalency and CpG methylation. A)** PCA analysis of ChIP-seq samples shows that biological replicates cluster together, while different histone PTMs (H3.3K27me3, H3K27me3, H3.3, and H3) cluster separately from each other. Furthermore, time points (day 0 and day 6) show separation as well. **B)** Based on RNA-seq data, genes were stratified into the quintiles based on their relative expression levels. The mean ChIP-seq signal for H3.3 and H3 is plotted across genes in the top quintile with the highest expression (HI, dark blue), the bottom quintile with the lowest expression (LO, yellow), and the middle three quintiles (MED, light blue), in 10 bp bins. The regions 3 kb upstream of the transcription start site (TSS) and 3 kb downstream of the transcription end site (TES) are also included. The inner tick marks demarcate 1 kb beyond the TSS and 1 kb before the TES. Inside this region, scaling has been applied to represent genes as a uniform length. Biological replicates are plotted separately, for both day 0 and day 6. The y-axis is CPM = counts per million mapped reads. **C)** H3.3K27me3 and H3K27me3 signal across CpG islands is plotted, and determines that H3K27me3 is more enriched at CpG islands, particularly at day 0. Heatmap is organized as described in panel B. **D)** The mean ChIP-seq signal for variant H3.3K27me3 or canonical H3K27me3 is plotted across 2251 previously identified bivalent genes[7] in 10 bp bins. The regions 3 kb upstream of the transcription start site (TSS) and 3 kb downstream of the transcription end site (TES) are also included. The inner tick marks demarcate 1 kb beyond the TSS and 1 kb before the TES. Inside this region, scaling has been applied to represent genes as a uniform length. Two biological replicates were analyzed. The y-axis is CPM = counts per million mapped reads.

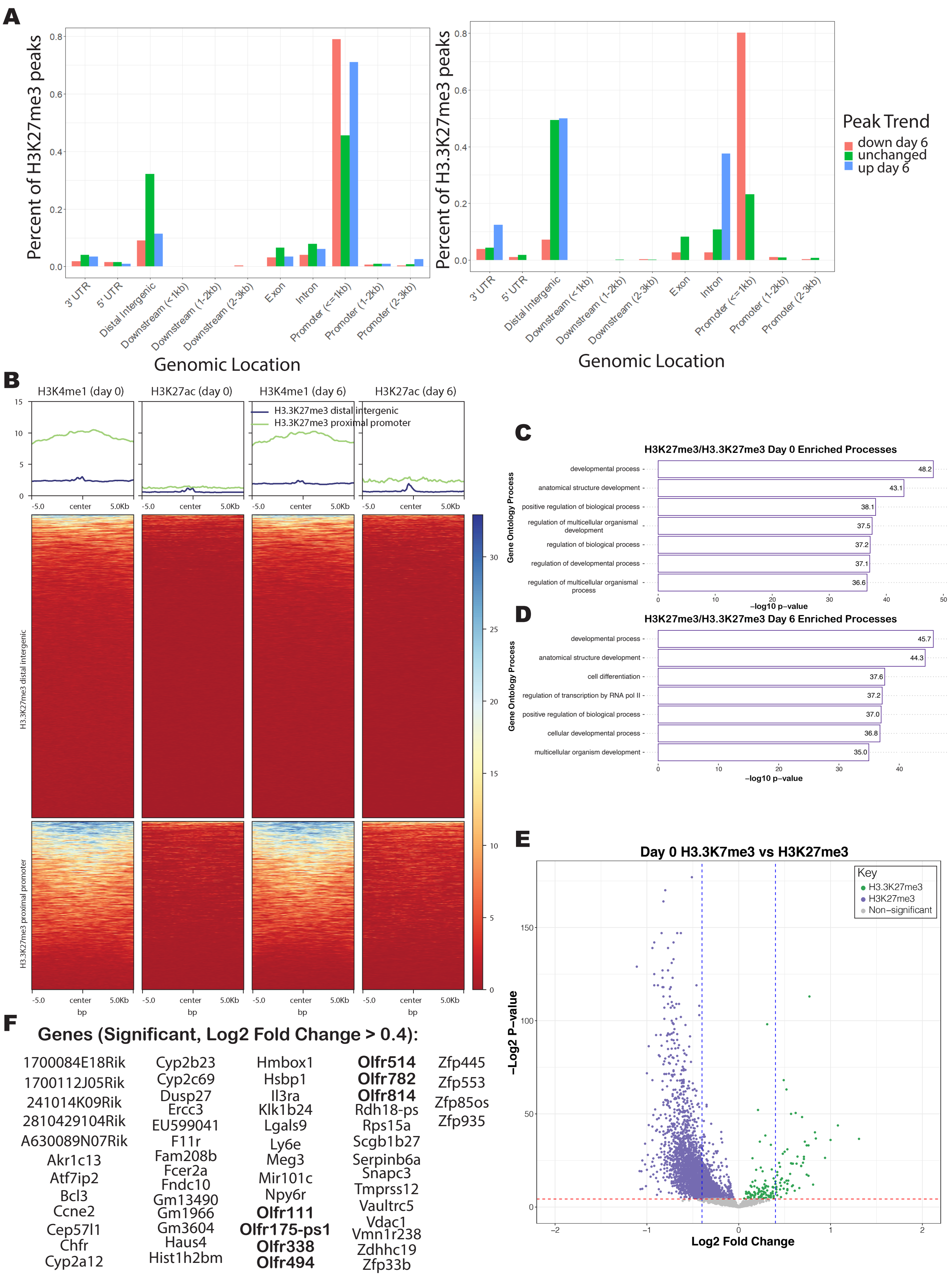
A
Percent of H3K27me3 peaks
Percent of H3.3K27me3 peaks
Genomic Location
3' UTR
5' UTR
Distal Intergenic
Downstream (<1kb)
Downstream (1-2kb)
Downstream (2-3kb)
Exon
Intron
Promoter (<=1kb)
Promoter (1-2kb)
Promoter (2-3kb)
Peak Trend
down day 6
unchanged
up day 6
B
H3K4me1 (day 0)
H3K27ac (day 0)
H3K4me1 (day 6)
H3K27ac (day 6)
H3.3K27me3 distal intergenic
H3.3K27me3 proximal promoter
-5.0
center
5.0Kb
bp
C
H3K27me3/H3.3K27me3 Day 0 Enriched Processes
Gene Ontology Process
developmental process 48.2
anatomical structure development 43.1
positive regulation of biological process 38.1
regulation of multicellular organismal development 37.5
regulation of biological process 37.2
regulation of developmental process 37.1
regulation of multicellular organismal process 36.6
−log10 p-value
D
H3K27me3/H3.3K27me3 Day 6 Enriched Processes
developmental process 45.7
anatomical structure development 44.3
cell differentiation 37.6
regulation of transcription by RNA pol II 37.2
positive regulation of biological process 37.0
cellular developmental process 36.8
multicellular organism development 35.0
−log10 p-value
E
Day 0 H3.3K7me3 vs H3K27me3
Key
H3.3K27me3
H3K27me3
Non−significant
−Log2 P−value
Log2 Fold Change
F
Genes (Significant, Log2 Fold Change > 0.4):
1700084E18Rik
1700112J05Rik
241014K09Rik
2810429104Rik
A630089N07Rik
Akr1c13
Atf7ip2
Bcl3
Ccne2
Cep57l1
Chfr
Cyp2a12
Cyp2b23
Cyp2c69
Dusp27
Ercc3
EU599041
F11r
Fam208b
Fcer2a
Fndc10
Gm13490
Gm1966
Gm3604
Haus4
Hist1h2bm
Hmbox1
Hsbp1
Il3ra
Klk1b24
Lgals9
Ly6e
Meg3
Mir101c
Npy6r
Olfr111
Olfr175-ps1
Olfr338
Olfr494
Olfr514
Olfr782
Olfr814
Rdh18-ps
Rps15a
Scgb1b27
Serpinb6a
Snapc3
Tmprss12
Vaultrc5
Vdac1
Vmn1r238
Zdhhc19
Zfp33b
Zfp445
Zfp553
Zfp85os
Zfp935

**Supplemental Figure 8. H3.3K27me3 shows unique genomic enrichment compared to H3K27me3, but does not overlap with enhancers. A)** Genomic localization counts for ChIP-seq peaks that went up at day 6 (blue), down on day 6 (red), or are unchanged between time points (green) for H3.3K27me3 (right plot) and H3K27me3 (left plot). The counts are graphed as a percent of the total number of peaks detected for that histone modification. Regulated expression criteria: adjusted p-value < 0.05 and log2 fold change greater than 2 standard deviations away from the mean of the distribution, i.e. z score > 2 or < -2. **B)** H3.3K27me3 (as annotated at distal intergenic regions or proximal promoter regions) ChIP-seq signal at both day 0 and day 6 was plotted across regions previously characterized to be marked by the enhancer marks H3K4me1 and H3K27ac in pluripotent (day 0) mESC [8]. The regions 5 kb upstream and 5 kb downstream of the peak region are also included. Overlap of H3.3K27me3 and H3K4me1 is only observed at proximal promoter regions. **C)** Gene ontology analysis was performed using GOrilla[9] on the ChIP-seq peaks more highly marked in H3K27me3 compared to H3.3K27me3 at day 0. **D)** Gene ontology analysis was performed using GOrilla[9] on the ChIP-seq peaks more highly marked in H3K27me3 compared to H3.3K27me3 at day 6. **E)** Volcano plot of the ChIP-seq data for day 0 (pluripotency) plotting the log2 fold change H3.3K27me3/H3K27me3 on the x-axis and the -log2 t-test p-value on the y-axis. The ChIP-seq data was filtered to only plot peaks annotated as being at promoter regions. The red dotted line indicates a p-value of 0.05 (-log2 p-value = 4.32). Anything above the red line is statistically significant. The blue dotted lines indicate a log2 fold change of -0.4 and +0.4. Green dots indicate peaks significantly enriched in the H3.3K27me3 ChIP-seq, and purple dots indicate peaks significantly enriched in the H3K27me3 ChIP-seq. The volcano plot is asymmetric because there are more regions highly marked in H3K27me3 than H3.3K27me3. **F)** List of the genes with a log2 fold change above 0.4 that are statistically significant as being more highly enriched in H3.3K27me3 at their promoter region at day 0 (from the volcano plot shown in panel E, green dots to the right of the blue dotted line). Olfactory receptor genes are bolded.

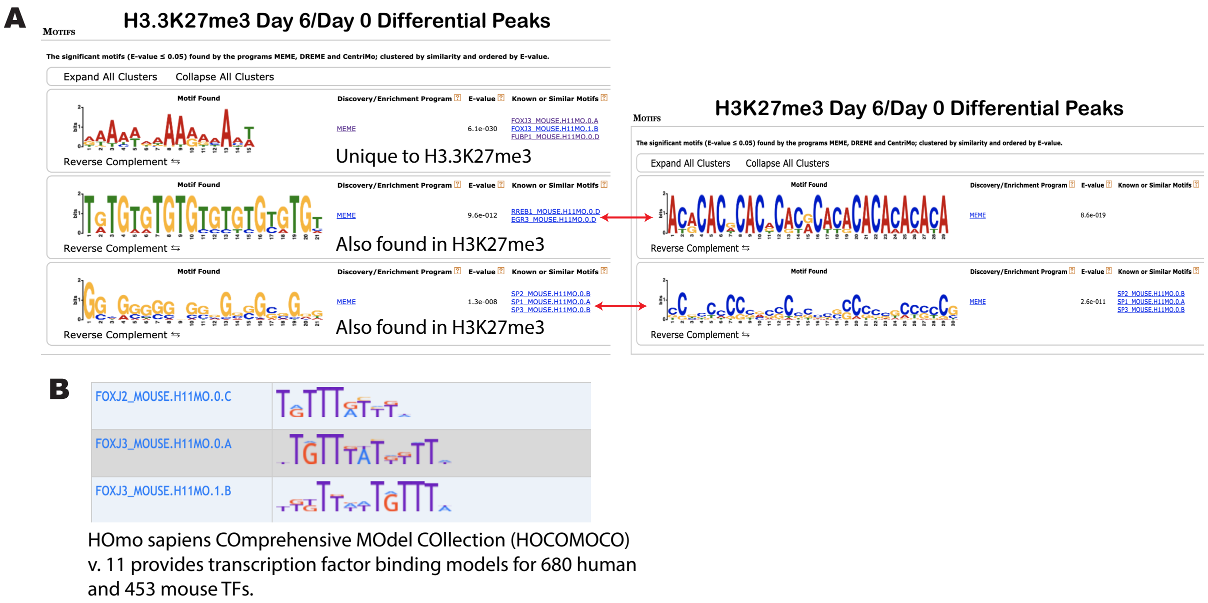


**Supplemental Figure 9. H3.3K27me3 differentiation peaks are uniquely enriched at FOXJ3 binding motifs during differentiation. A)** H3.3K27me3 peaks that were enriched at day 6 of differentiation compared to day 0 (pluripotency) and H3K27me3 peaks that were enriched at day 6 compared to day 0 were both analyzed by MEME-ChIP[10] and the motifs identified for each are shown in the snapshot of the results. The E-value is the "enrichment value" which is the significance value calculated by the MEME-ChIP program. The RREB1 and SP2 binding motifs were found to be significantly enriched for both H3.3K27me3 and H3K27me3, but the binding motif for FOXJ3 was only significantly enriched for H3.3K27me3. **B)** The FOXJ2 and FOXJ3 motifs in the HOmo sapiens COmprehensive MOdel COllection (HOCOMOCO) v. 11 database that the MEME-ChIP software uses for analysis of enriched binding motifs, which provides transcription factor binding models for 680 and human and 453 mouse transcription factors. The binding motifs for FOXJ2 and FOXJ3 show high sequence similarity.

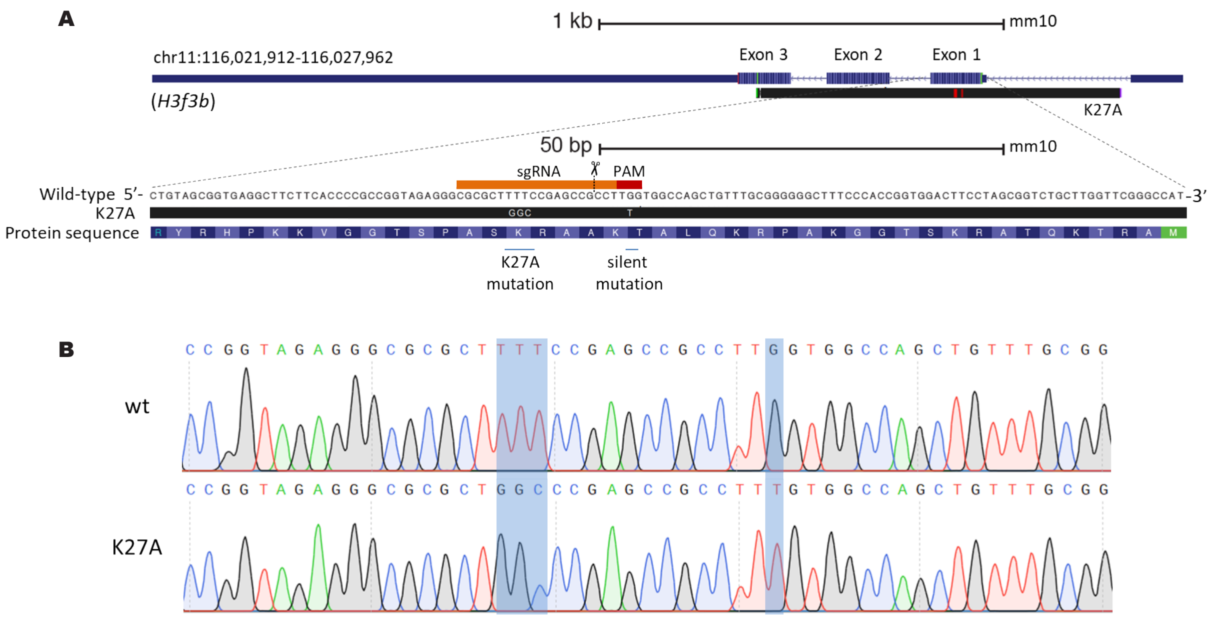


**Supplemental Figure 10. Generation and validation of the H3.3 K27A mutant. A)** A sgRNA (orange) was designed to target the first coding exon of the H3f3b gene, close to the lysine 27 codon sequence. Snapshots of the H3f3b locus from the UCSC Genome Browser [11] (http://genome.ucsc.edu/) are displayed, showing the alignment between the sequence amplified from the K27A mutant (black) and the wild-type mouse genome (GRCm38/mm10). In addition to the lysine-to-alanine substitution at position 27, one extra silent mutation (G > T) was introduced to disrupt the PAM sequence and hence prevent re-cleavage of the successfully edited locus. **B)** Inspection of the Sanger sequencing chromatograms confirms the introduction of the intended substitutions in homozygosity (highlighted).

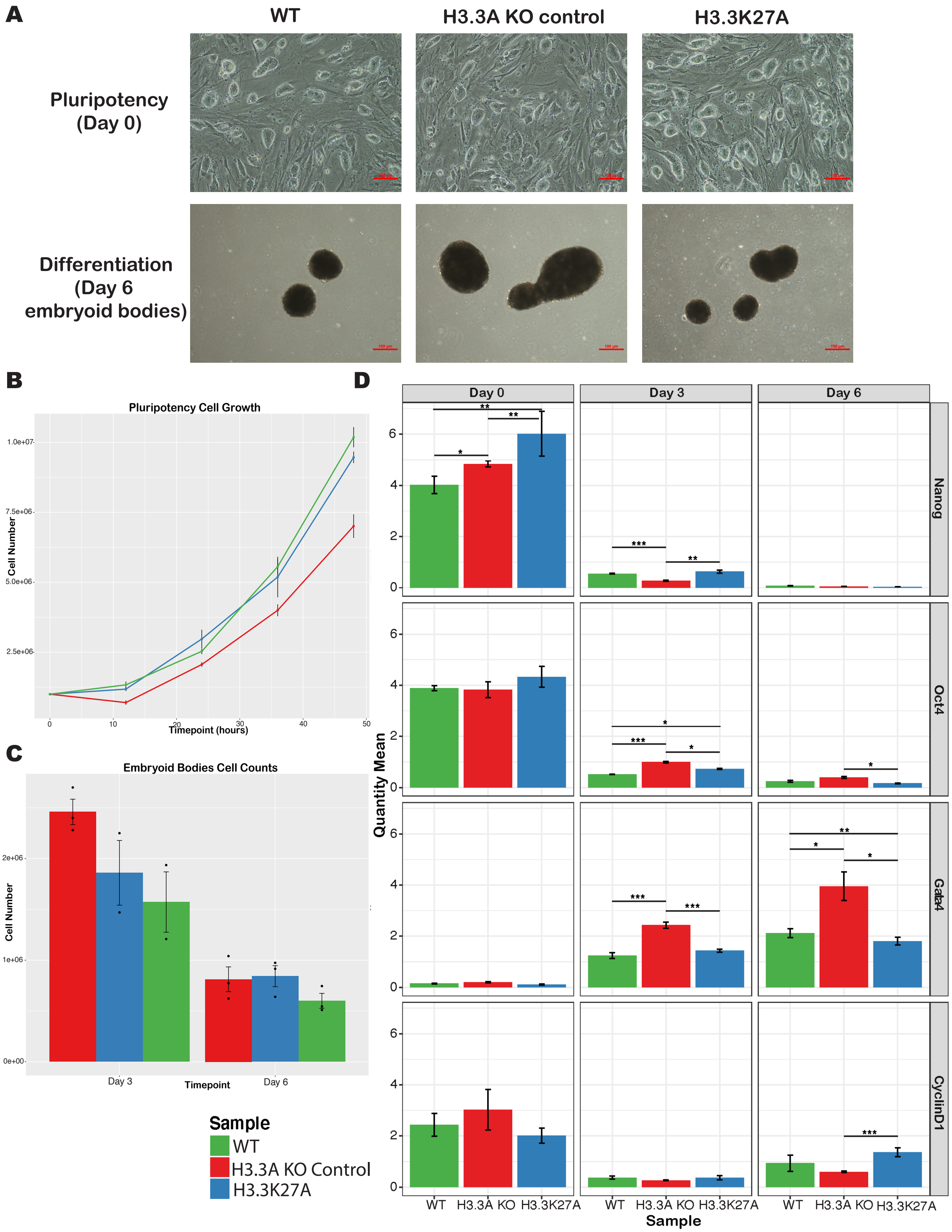

A
WT
H3.3A KO control
H3.3K27A
Pluripotency (Day 0)
Differentiation (Day 6 embryoid bodies)
B
Pluripotency Cell Growth
Cell Number
Timepoint (hours)
C
Embryoid Bodies Cell Counts
Cell Number
Day 3
Day 6
Timepoint
D
Day 0
Day 3
Day 6
Nanog
Oct4
Gata4
CyclinD1
Quantity Mean
WT
H3.3A KO
H3.3K27A
Sample
Sample
WT
H3.3A KO Control
H3.3K27A

**Supplemental Figure 11. No significant phenotypic differences between H3.3K27A mutants compared to controls, but some dysregulation of pluripotency and differentiation markers is detected. A)** Photographs of mESC grown in pluripotency on MEF feeder cells (top row) and in differentiation as embryoid bodies at the day 6 time point (bottom row) are shown for the wild-type (WT) cells, H3.3A knockout control cells (H3.3A KO control), and H3.3K27A mutants. Significant phenotypic differences not observed. **B)** A cell growth assay was conducted during pluripotency. Cell counts (y-axis) are plotted across the time points (x-axis) for WT (green), H3.3A KO control (red), and H3.3K27A (blue) cells. The mean of biological triplicates is plotted, with the standard deviation of the mean indicated. **C)** A cell growth assay was conducted during differentiation. Cell counts (y-axis) are plotted for day 3 and day 6 of differentiation for WT (green), H3.3A KO control (red), and H3.3K27A (blue) cells. The mean of biological replicates is plotted, with the standard deviation of the mean indicated. **D)** RT-PCR for pluripotency markers (Nanog and Oct4), and differentiation markers (Gata4 = endoderm marker, CyclinD1 = neuroectoderm marker) was conducted for WT (green), H3.3A KO control (red), and H3.3K27A (blue) cells across time points (day 0 = pluripotency, and day 3 and day 6 of differentiation). The quantity mean for each marker was normalized to a housekeeping gene, GAPDH. The normalized quantity mean is plotted, with standard deviation of the mean indicated. A Student's Type II T-test was used to calculate significance between samples. One asterisk indicates a p-value < 0.05, two asterisks indicate a p-value < 0.01, and three asterisks indicate a p-value < 0.001.

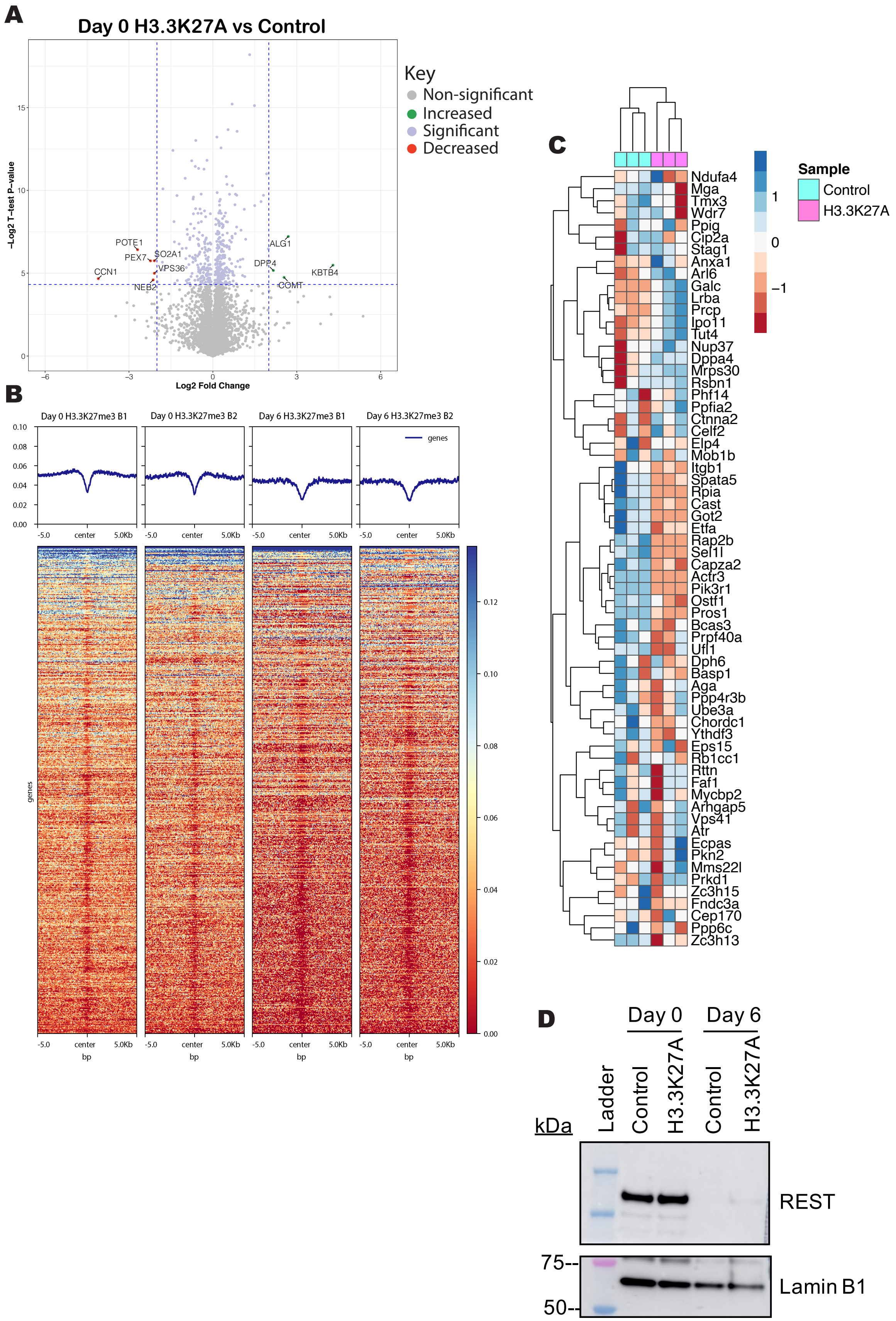
A
Day 0 H3.3K27A vs Control
Key
Non-significant
Increased
Significant
Decreased
-Log2 T-test P-value
Log2 Fold Change
POTE1
PEX7
SO2A1
VPS36
CCN1
NEB2
ALG1
DPP4
KBTB4
COMT
B
Day 0 H3.3K27me3 B1
Day 0 H3.3K27me3 B2
Day 6 H3.3K27me3 B1
Day 6 H3.3K27me3 B2
genes
-5.0
center
5.0Kb
bp
C
Sample
Control
H3.3K27A
Ndufa4
Mga
Tmx3
Wdr7
Ppig
Cip2a
Stag1
Anxa1
Arl6
Galc
Lrba
Prcp
Ipo11
Tut4
Nup37
Dppa4
Mrps30
Rsbn1
Phf14
Ppfia2
Ctnna2
Celf2
Elp4
Mob1b
Itgb1
Spata5
Rpia
Cast
Got2
Etfa
Rap2b
Sel1l
Capza2
Actr3
Pik3r1
Ostf1
Pros1
Bcas3
Prpf40a
Ufl1
Dph6
Basp1
Aga
Ppp4r3b
Ube3a
Chordc1
Ythdf3
Eps15
Rb1cc1
Rttn
Faf1
Mycbp2
Arhgap5
Vps41
Atr
Ecpas
Pkn2
Mms22l
Prkd1
Zc3h15
Fndc3a
Cep170
Ppp6c
Zc3h13
D
Day 0
Day 6
kDa
Ladder
Control
H3.3K27A
REST
Lamin B1
75--
50--

**Supplemental Figure 12. Proteomic dysregulation in H3.3K27A cells may not be directly due to loss of H3.3K27me3. A)** Volcano plot of proteomics data plotting the log2 fold change of H3.3K27A/control cells on day 0 (pluripotency) on the x-axis, and the -log2 t-test p-value is plotted on the y-axis. The horizontal blue line indicates a -log2 p-value of 4.32 (equivalent to a p-value of 0.05), so anything above that line is statistically significant. The vertical blue lines indicate a log2 fold change of +/- 2. Points indicated in green were significantly increased in the H3.3K27A mutant cells, above a log2 fold change of 2. Points indicated in red were significantly decreased in the H3.3K27A mutant (and thus increased in the control), below a log2 fold change of -2. Indicated in purple are all the other statistically significant proteins, regardless of a fold change cutoff. **B)** H3.3K27me3 ChIP-seq signal at both day 0 and day 6 was plotted across the TSS of genes of the dysregulated proteins identified in the H3.3K27A cells during differentiation. The regions 5 kb upstream and 5 kb downstream of the TSS are also included. Enrichment of H3.3K27me3 at these genes is only very slightly observed at day 0. **C)** Genes identified as being marked by H3.3K27me3 via ChIP-seq were detected as proteins in the H3.3K27A differentiation proteomics data, and heatmap analysis of the z-score indicates that upon loss of H3.3K27me3, as is the case in the H3.3K27A cells (pink), these proteins display variable expression compared to control cells (turquoise). **D)** Western blot analysis as presented in Figure 7E, but including day 0 (pluripotency). REST levels are high in pluripotency in both H3.3K27A and control cells, which drowns the signal from day 6 of differentiation. However, increased REST in H3.3K27A cells compared to control cells at day 6 is observed, although not to the same expression level as day 0. Lamin B1 is the loading control.

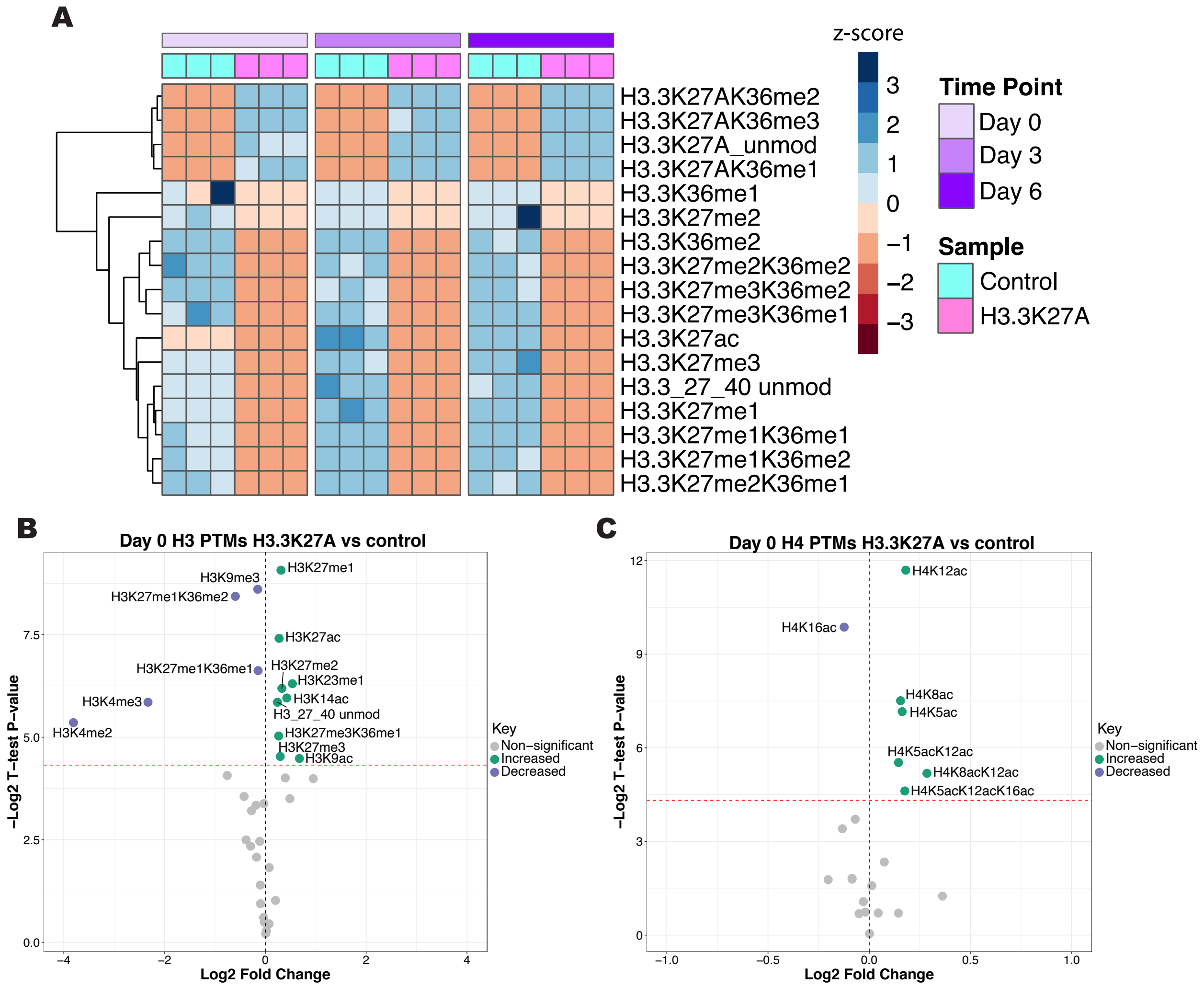


**Supplemental Figure 13. Detection of H3.3K27A peptide and loss of methylation at H3.3K27 is confirmed by MS analysis, resulting in slight global histone PTM changes at day 0. A)** The z-scores of the relative abundances of the detected histone H3.3 PTMs are plotted as a heatmap using hierarchal clustering. The heatmap is separated by time point: day 0 (pluripotency, indicated in light purple), day 3 of differentiation (purple), and day 6 of differentiation (dark purple). The three biological replicates for the H3.3K27A mutant cells is indicated in pink and the three biological replicates for the control cells is indicated in turquoise. **B)** A volcano plot of the H3 PTMs on day 0 (pluripotency). The log2 fold change of H3.3K27A/control is plotted with the -log2 t-test p-value on the y-axis. The red dotted line indicates a -log2 p-value of 4.32 (equivalent to a p-value of 0.05). Points indicated in green are significantly increased in the H3.3K27A cells compared to the control cells, and points indicated in purple are significantly decreased in the H3.3K27A cells compared to the control cells. **C)** A volcano plot of the H4 PTMs on day 0. The plot is organized in the same way as described for panel B.
